# Biased projection neuron–Kenyon cell connectivity shapes odor representations and learning in the *Drosophila* mushroom body

**DOI:** 10.1101/2025.10.29.684686

**Authors:** Alexander John MacKenzie, Lia Beatty, Andrew R. Butts, Jilian Mae Ulibarri, Dua Azhar, Prisca Amematsro, Sophie Jeanne Cécile Caron

## Abstract

Learning and memory centers must balance maximizing coding capacity with prioritizing biologically relevant information. Expansion layers, a circuit motif common to many learning and memory centers including the insect mushroom body, transform dense sensory representations into sparse, distributed ones, and theoretical models propose that random connectivity within these layers maximizes coding capacity by generating highly discriminable responses. Yet this solution creates a fundamental problem: purely random connectivity treats all stimuli equally, without prioritizing survival-relevant cues over neutral ones. The *Drosophila melanogaster* mushroom body has been shown to depart from pure randomness through systematic biases in projection neuron–Kenyon cell connectivity. Although connectivity appears random at the single-cell level, some projection neuron types connect up to 15-fold more frequently than others. Here, we find that these biases relate to function: the number of responding Kenyon cells tended to increase with projection neuron connectivity, and to correlate with learning performance. Odors activating more than 20% of Kenyon cells support more robust associative memories, whereas those activating fewer than 10% were poorly learned. VL1 projection neurons were a notable exception: despite their weak connectivity, they elicited broad Kenyon cell activity but failed to support learning, which pointed to a circuit-level gate on learning. These results suggest that connectivity biases in the mushroom body might prioritize ethologically relevant odors while preserving coding capacity for diverse associations.

## INTRODUCTION

Brains are built to learn. At any given moment, sensory systems flood the brain with information, but only a fraction of that information is used to learn and form memories (Xia et al., 2024). This selectivity reflects a fundamental trade-off that constrains learning and memory centers: these centers, with their limited number of neurons and synapses, must maintain the capacity to encode a wide range of stimuli, yet they must also reliably prioritize recurring, survival-relevant cues an animal is likely to encounter throughout its lifetime. This trade-off is expected to constrain the circuit architecture of learning and memory centers: architectures that maximize sensory coding capacity may not be optimal for selective encoding of highly relevant cues, and *vice versa* (Litwin-Kumar et al., 2017; Cayco-Gajic and Silver, 2019; Nguyen et al., 2023). Learning and memory centers must therefore be wired in ways that balance coding capacity with biological selectivity. How this balance is achieved remains poorly understood. Expansion layers are a well-characterized circuit motif found across diverse systems with different learning and memory functions that may provide insight into how circuit architectures balance capacity with selectivity.

Expansion layers are characterized by a small number of input neurons connecting to a relatively larger number of coding neurons, transforming sensory inputs into sparse, distributed representations (Babadi and Sompolinsky, 2014; Cayco-Gajic and Silver, 2019). This circuit motif appears in both vertebrates and invertebrates, and across diverse brain centers that support different forms of learning and memory. In the vertebrate cerebellum, a relatively small number of mossy fiber inputs connect to a greatly larger number of granule cells that generate sparse representations critical for motor learning and coordination (Marr, 1969; Albus, 1971; Cayco-Gajic and Silver, 2019). In invertebrates, similar expansion layers are found in the insect mushroom body, an associative center that transforms olfactory information into memories (Aso et al., 2014; Cognigni et al., 2018). Theoretical models propose that in these systems, stochastic or “random”, connectivity maximizes coding capacity by generating sparse representations and high discriminability (Babadi and Sompolinsky, 2014; Litwin-Kumar et al., 2017). Yet purely random connections treat all stimuli equally, failing to prioritize survival-relevant stimuli, thus maximizing capacity at the expense of selectivity.

Whether biological expansion layers deviate from pure randomness to resolve this trade-off can be addressed in the *Drosophila melanogaster* mushroom body, a numerically simple system comprising approximately 2,000 Kenyon cells that receive input from 51 olfactory glomeruli through projection neurons (Aso et al., 2014; Zheng et al., 2022; Grabe et al., 2015). Each Kenyon cell receives input from approximately six projection neurons through specialized dendritic structures termed “claws” (Butcher et al., 2012; Caron et al., 2013). While connectivity appears random at the single-cell level, systematic biases exist across the population: the identity of the projection neurons converging onto any individual Kenyon cell is stochastic, but the probability that a given projection neuron type is sampled differs systematically across types (Caron et al., 2013; Hayashi et al., 2022; Zheng et al., 2022; Ellis et al., 2024).

If the mushroom body optimally resolves the capacity-selectivity trade-off, these connectivity biases should represent an evolutionary compromise: sacrificing some coding capacity — which would be maximized by random, uniform connectivity — to selectively prioritize the representation of biologically relevant glomerular channels. This predicts that projection neurons with higher connectivity frequencies should process behaviorally critical odors and support stronger associative learning. We tested this hypothesis in three steps: first, we established that connectivity biases are stereotyped architectural features of the mushroom body; second, we determined whether connectivity biases correlate with Kenyon cell responses; and third, we examined whether connectivity biases are associated with learning performance.

## RESULTS

### Projection neuron–Kenyon cell connectivity shows systematic biases

Projection neurons connect individual olfactory glomeruli of the antennal lobe to the Kenyon cells of the mushroom body (Figure 1A-B). Connectivity biases have been reported across multiple datasets, with some projection neurons connecting to Kenyon cells far more frequently than others (Caron et al., 2013; Hayashi et al., 2022; Zheng et al., 2022; Ellis et al., 2024). For example, DL3 projection neurons connect to Kenyon cells less frequently than expected under a uniform distribution, while DA1 projection neurons connect more frequently (Figure 1C-D). Whether these biases are stereotypical architectural features of the *Drosophila* mushroom body or dataset specific observations has not been established. We therefore quantified their magnitude and reproducibility across multiple independent datasets.

**Figure 1.**
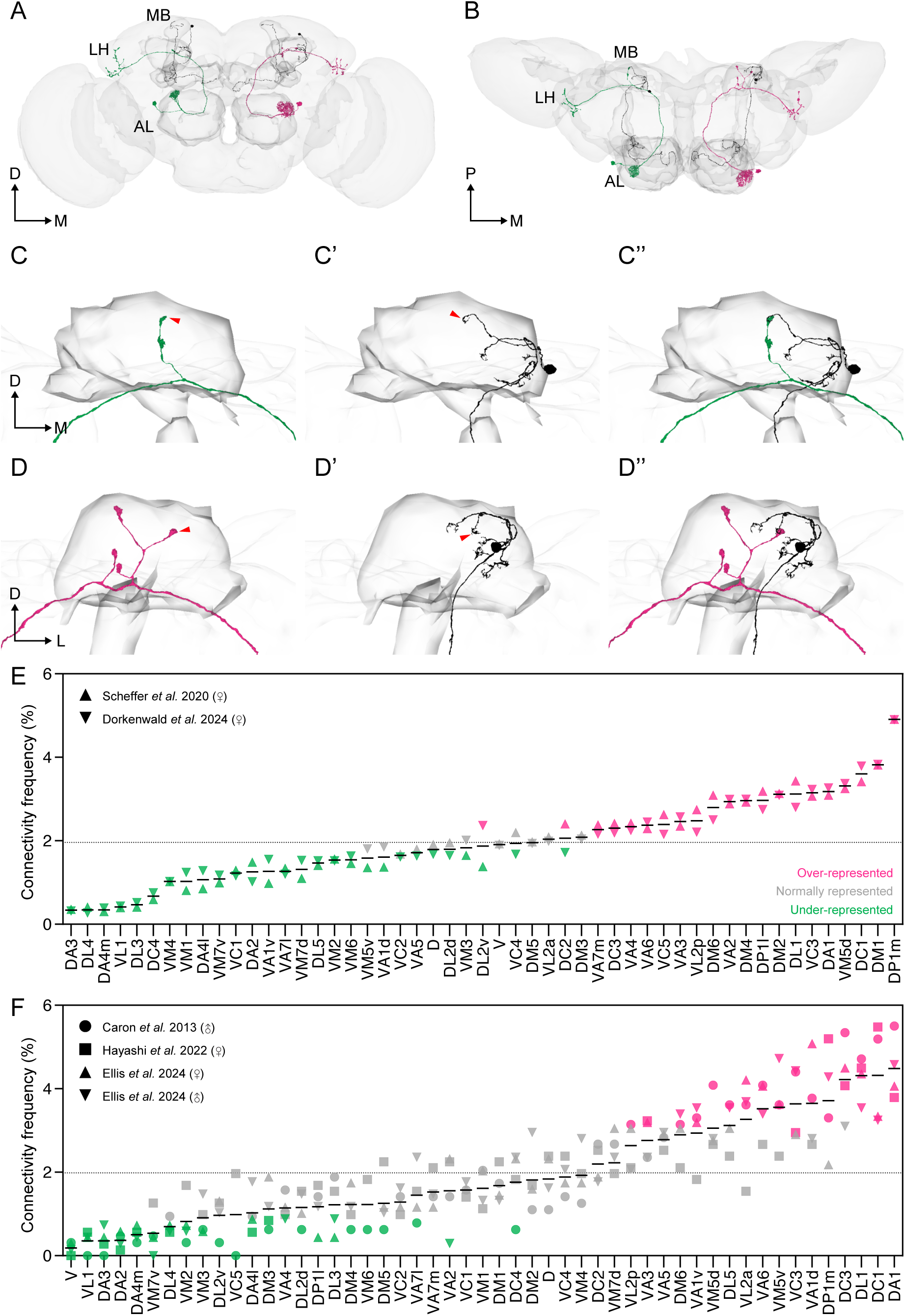
Projection neuron–Kenyon cell connections are systematically biased. (A,B) Electron microscopy–based reconstructions of DL3 (green) and DA1 (pink) projection neurons innervating glomeruli in the antennal lobe (AL). Each projection neuron forms *en passant* synapses with Kenyon cells (black) in the mushroom body (MB) and terminates in the lateral horn (LH). Panels show anterior (A) and dorsal (B) views. Outlines of the antennal lobe and mushroom body are included for anatomical context (gray). Orientation arrows indicate dorsal (D), medial (M), and posterior (P) directions. (C) A Kenyon cell (black) forms five claws, one of them (red arrowhead) receiving input from a DL3 projection neuron (green). (D) A Kenyon cell (black) forms six claws, two of them (red arrowheads) receiving input from a DA1 projection neuron (pink). (E) Distribution of glomerular input frequencies to Kenyon cells in two electron microscopy–based connectomes. Each data point represents the proportion of synapses contributed by projection neurons from a given glomerulus to the entire Kenyon cell population. The dotted line indicates the connectivity frequency expected under uniform connectivity (1.96%, that is, 1/51 glomeruli). Symbols indicate datasets: upward triangle (Scheffer et al., 2020), downward triangle (Dorkenwald et al., 2024). Colors indicate degree of representation: pink (overrepresented glomeruli), green (underrepresented glomeruli), gray (not significantly biased glomeruli). (F) Distribution of glomerular input frequencies in four large-scale tracing datasets. The dotted line indicates the connectivity frequency expected under uniform connectivity (1.96%, that is, 1/51 glomeruli). Symbols indicate datasets: circle (Caron et al., 2013), square (Hayashi et al., 2022), upward triangle (Ellis et al., 2024, female), downward triangle (Ellis et al., 2024, male). Colors as in E. See Table S1.

We first analyzed two complete connectomes that reconstructed full mushroom bodies: the hemibrain connectome and the FlyWire whole brain connectome (Scheffer et al., 2020; Dorkenwald et al., 2024). These datasets report the number of synapses contributed by the projection neurons associated with each glomerulus to the entire Kenyon cell population. If connections were purely random, each of the 51 projection neuron types, corresponding to the 51 glomeruli, would contribute on average 2% of the total input, although discrete sampling produces deviations from this mean even under random connectivity. Instead, and consistent with previous analyses of these same datasets (Li et al., 2020; Zheng et al., 2022), we observed a markedly non-uniform distribution, with individual glomeruli contributing between 0.34% and 4.91% of total inputs, about 15-fold difference, indicating substantial biases (Table S1). The identities of over- and underrepresented glomeruli were consistent across both connectomes (Figure 1E).

We then analyzed four large-scale connectivity matrices generated from dye-filling and single-cell tracing experiments (Caron et al., 2013; Hayashi et al., 2022; Ellis et al., 2024). In these datasets, projection neuron inputs to individual Kenyon cells were systematically traced, with each matrix comprising at least 200 Kenyon cells. Unlike the electron microscopy–based connectomes, these matrices reflect the frequency with which projection neurons from specific glomeruli connect to Kenyon cells, independent of synaptic weight. These datasets also revealed a non-uniform distribution of inputs, with glomerular contributions ranging from 0.19% to 4.48% (Table S1). Only twelve glomeruli were consistently represented at near-expected levels in all four matrices (Figure 1F). The identities of the most strongly over- and underrepresented glomeruli were largely consistent across the electron microscopy–based connectomes and large-scale tracing datasets, although some glomeruli differed in their ranking between methods.

Together, these findings confirm that projection neuron–Kenyon cell connectivity is systematically biased and that the same projection neurons are over- or underrepresented across datasets.

### Bouton morphology predicts connectivity biases

Connectivity biases have been attributed to the number of synaptic boutons collectively formed by the projection neurons associated with a given glomerulus in the calyx of the mushroom body, rather than to the number of projection neurons associated with that glomerulus (Zheng et al., 2022; Ellis et al., 2024). We sought to test this relationship using light microscopy. To visualize projection neurons with varying degrees of connectivity, we used publicly available *split-GAL4* transgenic lines (Shuai et al., 2025). Such lines are currently available for a limited number of projection neuron types — specifically the underrepresented VL1, DL3, and VM4 glomeruli and the overrepresented DA1 glomerulus (Figure 2A-D,G-J). Additional lines target projection neurons innervating pairs or trios of glomeruli, including the DA4l and DC3 glomeruli, or the VM5v, VM7d, and VM7v glomeruli (Figure 2E-F,K-L). Based on the combined connectivity frequencies of the glomeruli they target, both lines are expressed in projection neurons with a cumulative representation that exceeds that of the average glomerulus and are therefore classified as overrepresented.

**Figure 2.**
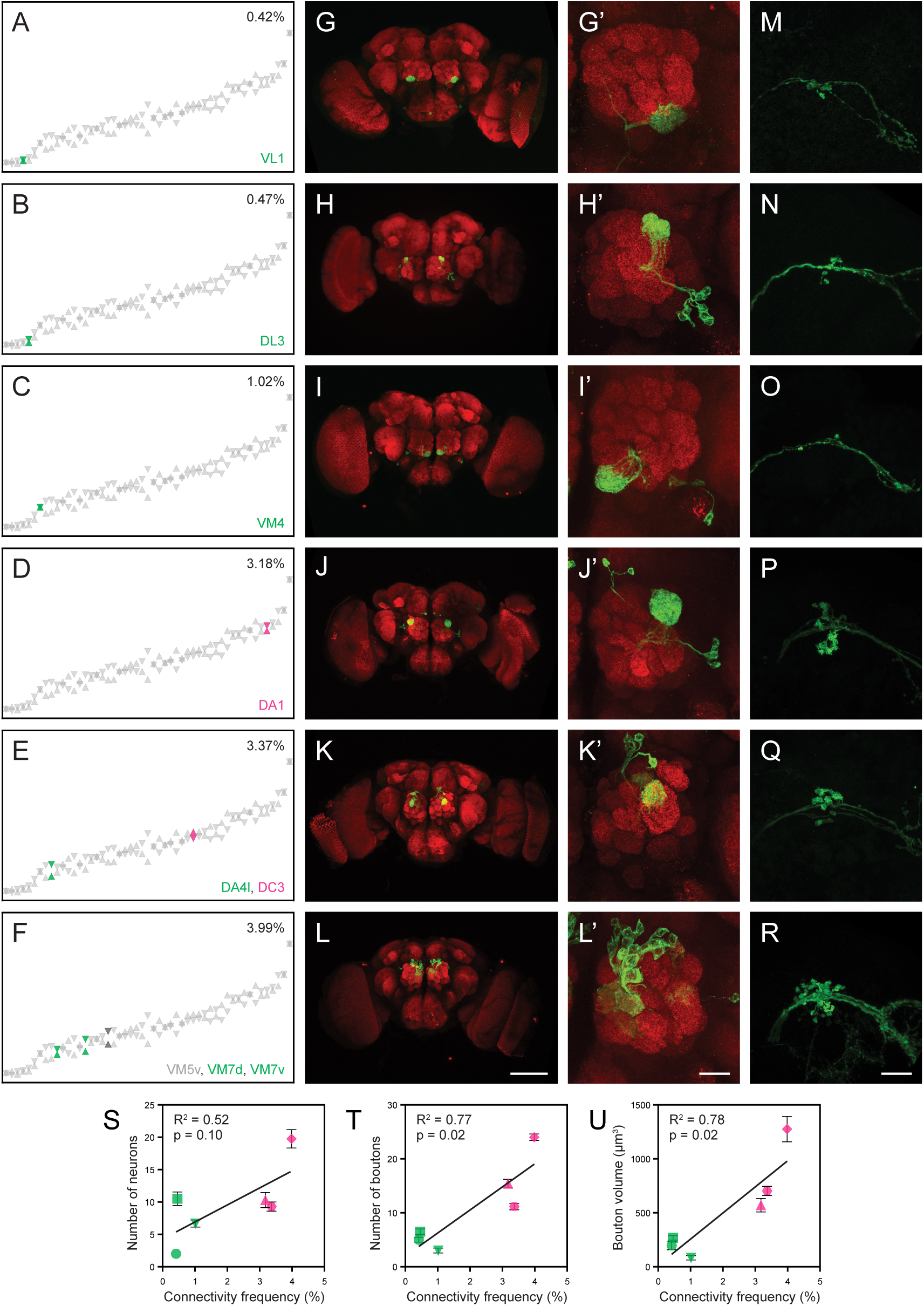
Morphological features of projection neurons correlate with connectivity frequency. (A–F) Distribution of connectivity frequencies from Figure 1E, with colored markers highlighting the six projection neuron types targeted by the *split-GAL4* lines used in this study. Lines target either individual glomeruli (VL1, DL3, VM4, DA1) or glomerular combinations (DA4l, DC3; VM5d, VM7d, VM7v). Numbers indicate average connectivity frequencies for lines targeting a single glomerulus or cumulative connectivity frequencies for lines targeting multiple glomeruli. Colors indicate degree of representation: pink (overrepresented glomeruli) or green (underrepresented glomeruli). (G–R) Expression patterns of projection neurons labeled by the *split-GAL4 lines*, visualized with *UAS-GFP* and immunostained for GFP (green) and Bruchpilot (magenta). Scale bar: 100 μm. (G′–L′) Higher magnification views of antennal lobe glomeruli showing projection neuron arborizations and somata. Scale bar: 20 μm. (M–R) Higher magnification views of mushroom body calyx showing presynaptic boutons. Scale bar: 20 μm. (S–U) Quantification of morphological features versus average connectivity frequency from electron microscopy-based connectomes. (S) Number of projection neurons *per* line. (T) Total bouton number in the calyx. (U) Total bouton volume. All values are reported per hemisphere. Each data point represents a *split-GAL4* line; error bars show standard error of the mean; linear regression lines are shown with R² and p values. Symbols indicate the identity of the *split-GAL4* lines: circle (VL1), square (DL3), downward triangle (VM4), upward triangle (DA1), hexagon (DA4l+DC3), diamond (VM5d+VM7d+VM7v). Colors indicate degree of representation: pink (overrepresented glomeruli) or green (underrepresented glomeruli). See Table 1 and Figure S1.

**Table 1.**
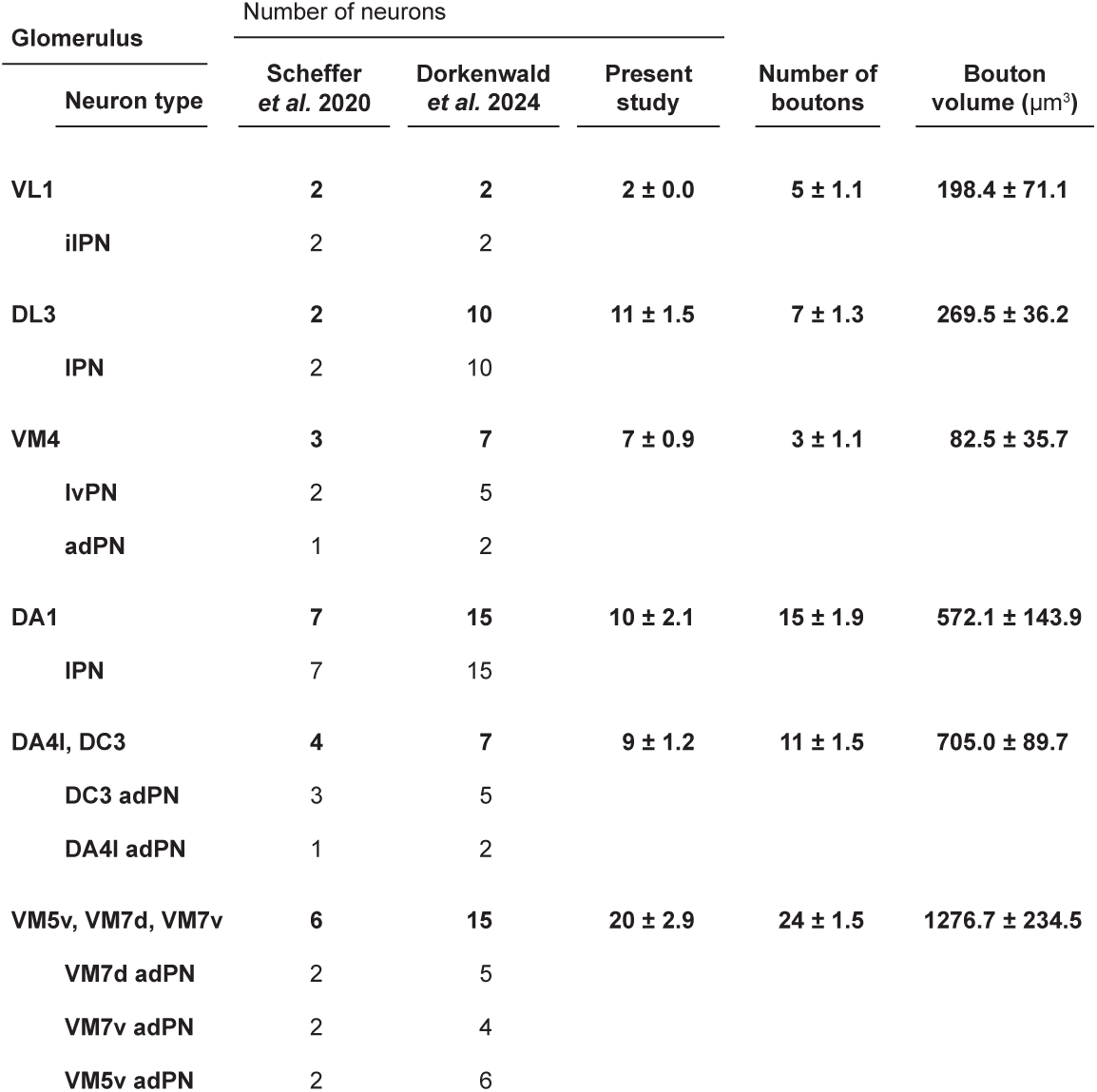
Morphological features of projection neurons labeled by *split-GAL4* lines. The table reports projection neurons expressing each *split-GAL4* transgene, their associated glomeruli, and quantified morphological features including number of neurons, number of boutons, and total bouton volume, all reported *per* hemisphere. For comparison, projection neuron counts for each glomerulus from two electron microscopy connectomes are included (Scheffer et al., 2020; Dorkenwald et al., 2024). Values are reported as means with standard error of the mean.

We used these transgenic lines in combination with a *UAS-GFP* reporter to quantify three morphological features of these projection neurons: the number of GFP-labeled neurons, the total number of boutons formed in the mushroom body calyx by all labeled neurons, and the total bouton volume (Table 1). All measurements are reported per hemisphere (Figure 2S-U, Table 1). We then compared each of these features to the average connectivity frequency of the corresponding glomeruli, based on data from both the electron microscopy–based connectomes and the large-scale tracing datasets (Figure 2S-U, Figure S1). Bouton number and bouton volume correlated with connectivity frequency (*R*² = 0.77, *p* = 0.02 and R² = 0.78, *p* = 0.02, respectively; Figure 2T,U). In general, projection neurons that formed more or larger boutons were more frequently connected to Kenyon cells. For example, projection neurons from the VL1 glomerulus — consistently underrepresented across datasets — had the fewest boutons and the smallest total bouton volume (Figure 2A,G,M,T,U). By contrast, projection neurons from the DA1 glomerulus had the highest bouton number and volume and were among the most highly represented (Figure 2D,J,P,T,U). The line expressing in the VM5v, VM7d, and VM7v projection neurons formed the largest number of boutons, consistent with the combined connectivity frequencies of these glomeruli (Figure 2F,L,R,T,U).

By comparison, the number of projection neurons per glomerulus correlated weakly with connectivity frequency (*R*² = 0.52, *p* = 0.10; Figure 2S). This was most evident with the DL3 glomerulus. Among lines restricted to a single glomerulus, DL3 had the highest number of projection neurons, with 11 ± 2 labeled neurons, yet it was underrepresented in both connectomes and in two of the four connectivity matrices (Table 1, Figure 2B,H,N,S). DL3 projection neurons formed few boutons relative to their number: many of their axonal tracts passed through the calyx without forming boutons. We note that DL3 projection neurons are absent from one of the two connectomes, either because they were not reconstructed or because they were not present in the reconstructed brain, and the DL3 projection neuron count in the other connectome is low relative to other glomeruli.

Together, these findings indicate that bouton number and volume correspond more closely to connectivity frequency than projection neuron number does.

### Connectivity frequency corresponds to the number of responding Kenyon cells, with VL1 as an exception

We next tested whether projection neuron–Kenyon cell connectivity biases correspond to differences in Kenyon cell responses. To address this, we developed an imaging approach that combined optogenetic stimulation of projection neurons with calcium imaging in Kenyon cells. Projection neurons that expressed the light-gated channel Chrimson under the control of *split-GAL4* transgenic lines were activated using 637 nm light, while Kenyon cells expressing GCaMP6f under LexA control were imaged following a brief light pulse across multiple optical planes using two-photon microscopy. The combination of these *split-GAL4* and *LexA* tools enabled us to specifically activate a given type of projection neuron while recording across Kenyon cells.

For each Kenyon cell, response magnitude was quantified as ΔF/F, and cells were initially categorized as either responders or non-responders (Figure 3A-L, Figure S2). The distribution of response amplitudes among responders was unimodal with a pronounced positive skew (Figure S3A-F). Based on this distribution, responders were further classified as weak or strong using k-means clustering on the maximum ΔF/F of the trial-averaged trace (Figure 3A’’-F’’, Figure S2C,D). We quantified three parameters for each sample: the percentage of responding Kenyon cells (“responders”), the frequency of weak and strong responders and the mean maximum peak amplitude among all responders. These parameters were then compared to the average connectivity frequency of the glomeruli stimulated in each experiment, as derived from electron microscopy–based connectomes or large-scale tracing datasets (Figure 3M-P, Figures S4A-D).

**Figure 3.**
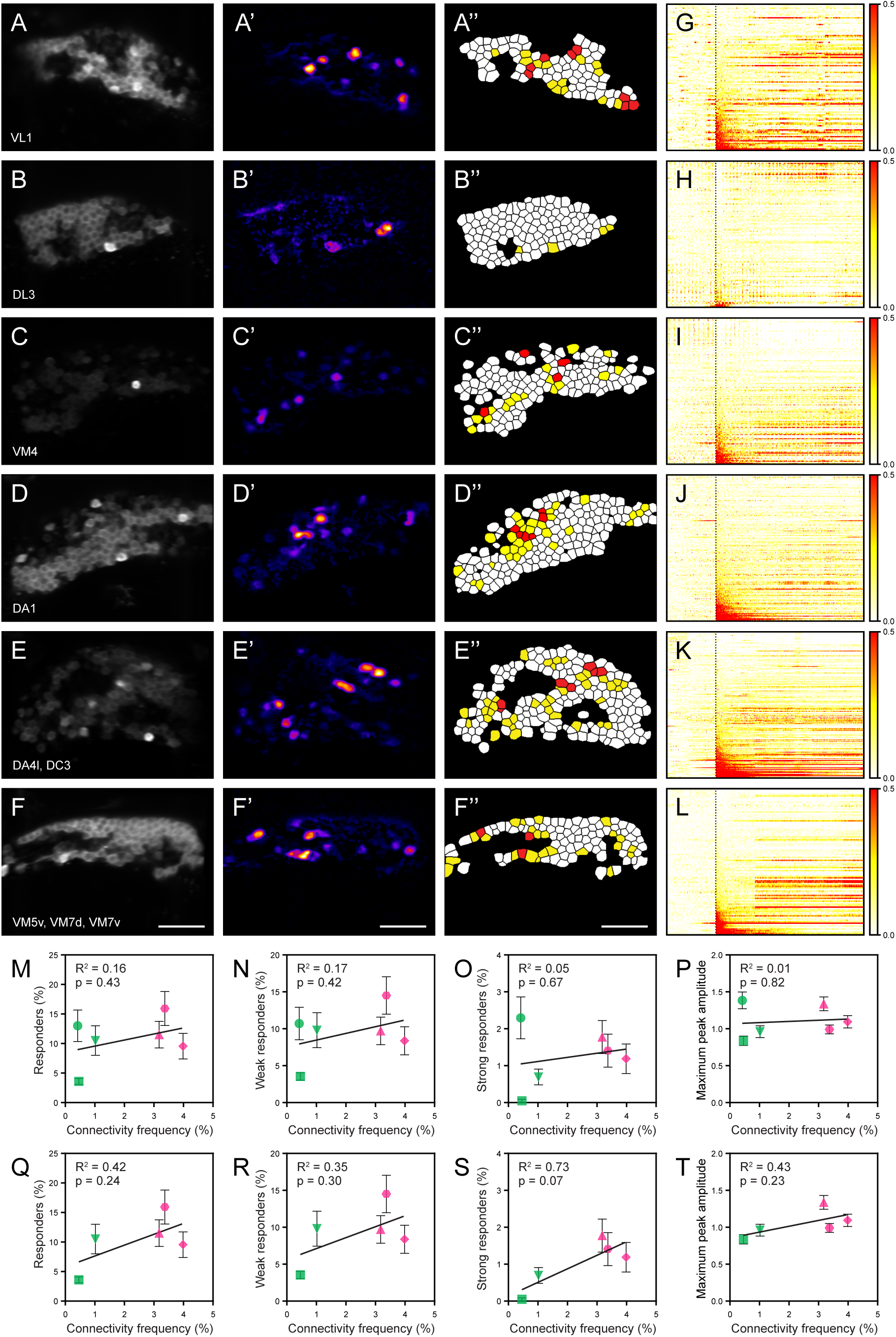
Connectivity frequency corresponds to the number of responding Kenyon cells. (A–F) Optogenetic activation of projection neurons from *split-GAL4* lines and corresponding responses in Kenyon cells. For each line, panels show: a representative fluorescence image of Kenyon cells expressing GCaMP6f in a single optical plane (A–F); a heat map of ΔF responses following light stimulation (A′–F′); and segmented activation map in which non-responsive Kenyon cells are filled in white, weak responders are filled in yellow, and strong responders are filled in red (A″–F″). Scale bar: 20 μm. (G–L) Representative raster plots showing calcium responses from all recorded Kenyon cells following optogenetic stimulation obtained using each of the *split-GAL4* lines: VL1 (G), DL3 (H), VM4 (I), DA1 (J), DA4l+DC3 (K), and VM5d+VM7d+VM7v (L). Each row represents one cell, each column represents one imaging frame, and color intensity indicates ΔF/F amplitude at that frame (scale: 0 to 0.5 standard deviation). The dotted vertical line marks stimulus onset. (M–T) Quantification of Kenyon cell responses plotted against connectivity frequency of stimulated projection neurons (averaged across both connectomes; Table 1). Panels M–P include all data; panels Q–T show the same analyses with VL1 excluded. (M,Q) Percentage of all responders. (N,R) Percentage of weak responders. (O,S) Percentage of strong responders. (P,T) Maximum peak ΔF/F amplitude. Each data point represents a *split-GAL4* line; error bars show standard error of the mean; linear regression lines are shown with R² and p values. Symbols indicate projection neuron identity: circle (VL1), square (DL3), downward triangle (VM4), upward triangle (DA1), hexagon (DA4l+DC3), diamond (VM5d+VM7d+VM7v). Colors indicate glomerular representation: pink (overrepresented) or green (underrepresented). See Table S2 and Figures S2–S13.

The percentage of weak and strong responders varied across *split-GAL4* lines and increased with the connectivity frequency of the stimulated projection neurons, although this correlation did not reach statistical significance (Figure 3M-O, Figure S4A-C, Figures S5-S11). The proportion of strong responders was higher when stimulating projection neurons with higher connectivity frequency — for example, in lines targeting the DA1 glomerulus or multiple glomeruli (Table S2, Figures S8-S11). In contrast, strong responders were nearly absent in lines targeting projection neurons with lower connectivity frequency, with VL1 being the only exception: its stimulation generated the largest number of strong responders (Table S2, Figure S5, Figure S11). When VL1 was excluded from the analysis, the correlation between connectivity frequency and strong responders was stronger but still did not reach significance (*R²* = 0.73, *p* = 0.07; Figure 3S). The mean maximum peak amplitude of responders did not correlate with connectivity frequency, either with or without VL1 included in the analysis, which is consistent with connectivity frequency corresponding to response likelihood rather than to response magnitude once a cell is responsive (Figure 3P,T, Figure S4D,H).

To explore potential mechanisms underlying the VL1 exception, we considered two possibilities. First, VL1 projection neurons may preferentially converge onto the same Kenyon cells, increasing the likelihood of coincident input. Our analysis of the FlyWire connectome provided no support for this hypothesis: only two Kenyon cells received two inputs from the VL1 glomerulus, a number consistent with random sampling (Figure S12A-B). Second, individual VL1 projection neuron–Kenyon cell connections may contain an unusually high number of synapses, allowing a single input to drive Kenyon cell response more effectively. This was also not supported: VL1 projection neurons had the second-lowest synapse count per Kenyon cell connection among all projection neurons (Figure S12C). A previous study reported that the VL1 glomerulus is associated with both excitatory and inhibitory projection neurons, with much of the inhibitory output directed to the lateral horn rather than the mushroom body (Bates et al. 2020a). The circuit basis for the broad Kenyon cell responses evoked by VL1 stimulation is therefore unresolved, and our data do not establish whether it involves direct excitation, disinhibition, or another mechanism.

Together, these results indicate that connectivity biases correspond to differences in the number of responding Kenyon cells.

### Odor-evoked Kenyon cell responses correspond to connectivity biases

Our optogenetic experiments indicated that the frequency with which projection neurons connect to Kenyon cells corresponds to the number of Kenyon cells that respond to their stimulation, suggesting that biased connectivity might shape odor-evoked responses. We therefore sought to identify odors that activate Kenyon cells at varying frequencies.

Odors differ in the number and identity of olfactory sensory neurons, and consequently, the glomeruli they activate (Hallem and Carlson, 2006; Wilson, 2013): some odors activate multiple glomeruli, which we refer to as “multi-glomerular odors”, while others primarily activate a single glomerulus, which we refer to as “mono-glomerular odors”. We hypothesized that multi-glomerular odors would activate more Kenyon cells than mono-glomerular odors. To test this, we imaged Kenyon cell responses to six odors with different glomerular activity patterns. Three mono-glomerular odors — geosmin (DA2), farnesol (DC3), and valencene (DC1) — were compared to three multi-glomerular odors — pyrrolidine (two glomeruli), 4-methylcyclohexanol (three glomeruli), and 3-octanol (ten glomeruli) (Table 2). Glomerular activation patterns were predicted from a publicly available database of olfactory responses (Münch and Galizia, 2016).

**Table 2.** Predicted glomerular activation and connectivity frequencies for tested odors. The table reports the predicted olfactory receptor and glomerulus activated by each of the tested odors, based on publicly available response data (Münch & Galizia, 2016); these predictions are derived at the receptor level, do not account for the concentrations used here, and may not reflect activation in the antennal lobe under our conditions. Mean connectivity frequencies of the corresponding projection neurons are provided from electron microscopy connectomes and large-scale tracing datasets (Caron et al., 2013; Li et al., 2020; Scheffer 2020; Hayashi et al., 2022; Dorkenwald et al., 2024). For multi-glomerular odors, cumulative connectivity frequency is the sum of the connectivity frequencies of the individual activated glomeruli.

| Odor |  | Connectivity frequency (%) |  |  |  | Connectivity frequency (%) |  |  |  |  |  |
| --- | --- | --- | --- | --- | --- | --- | --- | --- | --- | --- | --- |
|  |  | Scheffer<br>*et al.* 2020 | Dorkenwald<br>*et al.* 2024 | Average<br>connectivity | Cumulative<br>connectivity | Caron<br>*et al.* 2013 | Hayashi<br>*et al.* 2022 | Ellis *et al.*<br>2024 (♀) | Ellis *et al.*<br>2024 (♂) | Average<br>connectivity | Cumulative<br>connectivity |
| farnesol |  |  |  |  | 2.30 |  |  |  |  |  | 4.26 |
| DC3 | Or83c (0.922) | 2.41 | 2.19 | 2.30 |  | 5.35 | 4.07 | 4.50 | 3.10 | 4.26 |  |
| valencene |  |  |  |  | 3.60 |  |  |  |  |  | 4.31 |
| DC1 | Or19a (0.775) | 3.41 | 3.79 | 3.60 |  | 5.19 | 5.48 | 3.34 | 3.25 | 4.31 |  |
| geosmin |  |  |  |  | 1.25 |  |  |  |  |  | 0.37 |
| DA2 | Or56a (0.573) | 1.49 | 1.02 | 1.25 |  | 0.47 | 0.14 | 0.58 | 0.30 | 0.37 |  |
| 4-methylcyclohexanol |  |  |  |  | 7.37 |  |  |  |  |  | 6.48 |
| VA3 | Or67b (0.876) | 2.35 | 2.58 | 2.46 |  | 2.36 | 3.23 | 3.19 | 2.36 | 2.79 |  |
| D | Or69a (0.635) | 1.91 | 1.68 | 1.79 |  | 1.10 | 2.39 | 2.32 | 1.62 | 1.86 |  |
| DM2 | Or22a (0.254) | 3.13 | 3.09 | 3.11 |  | 1.10 | 1.69 | 1.60 | 2.95 | 1.83 |  |
| pyrrolidine |  |  |  |  | 3.73 |  |  |  |  |  | 3.44 |
| VL1 | Ir75d (0.973) | 0.40 | 0.44 | 0.42 |  | 0.00 | 0.56 | 0.44 | 0.44 | 0.36 |  |
| VM5d | Or85b (0.309) | 3.25 | 3.37 | 3.31 |  | 4.09 | 2.67 | 2.76 | 2.81 | 3.08 |  |
| 3-octanol |  |  |  |  | 25.84 |  |  |  |  |  | 25.77 |
| D | Or69a (0.741) | 1.91 | 1.68 | 1.79 |  | 1.10 | 2.39 | 2.32 | 1.62 | 1.86 |  |
| VM5d | Or85b (0.676) | 3.25 | 3.37 | 3.31 |  | 4.09 | 2.67 | 2.76 | 2.81 | 3.08 |  |
| VM5v | Or98a (0.592) | 1.36 | 1.81 | 1.58 |  | 3.62 | 2.39 | 3.63 | 4.73 | 3.59 |  |
| DC2 | Or13a (0.579) | 2.40 | 1.72 | 2.06 |  | 2.67 | 2.53 | 1.89 | 1.77 | 2.22 |  |
| DM3 | Or47a (0.468) | 2.13 | 2.04 | 2.09 |  | 0.63 | 0.84 | 1.89 | 1.18 | 1.14 |  |
| VC3 | Or35a (0.434) | 3.07 | 3.23 | 3.15 |  | 4.40 | 2.95 | 2.90 | 4.43 | 3.67 |  |
| DC1 | Or19a (0.374) | 3.41 | 3.79 | 3.60 |  | 5.19 | 5.48 | 3.34 | 3.25 | 4.31 |  |
| VA4 | Or85d (0.369) | 2.42 | 3.25 | 2.34 |  | 1.57 | 0.98 | 1.16 | 0.89 | 1.15 |  |
| DM2 | Or22a (0.337) | 3.13 | 3.09 | 3.11 |  | 1.10 | 1.69 | 1.60 | 2.95 | 1.83 |  |
| DM6 | Or67a (0.319) | 3.09 | 2.50 | 2.80 |  | 3.14 | 2.11 | 3.05 | 3.40 | 2.92 |  |

Kenyon cells expressing GCaMP6f under LexA control were imaged using two-photon microscopy during a three-second odor pulse delivered via an olfactometer (Figure 4A-L). For each sample, Kenyon cell somata were segmented, and odor-evoked calcium responses were analyzed following a procedure similar to that described in the previous section: based on response amplitude, each Kenyon cell was classified as a non-responder, weak responder or strong responder (Figures 4A’’-F’’, Figure S14). We quantified the same three response features as in the optogenetic experiments: the percentage of responders, the frequency of weak and strong responders, and the mean maximum peak amplitude among responders. These parameters were then compared to the average connectivity frequency of the glomeruli activated by each odor, as determined from either electron microscopy–based connectomes or large-scale projection neuron tracing datasets (Figures 4M-P, Figure S15A-D).

**Figure 4.**
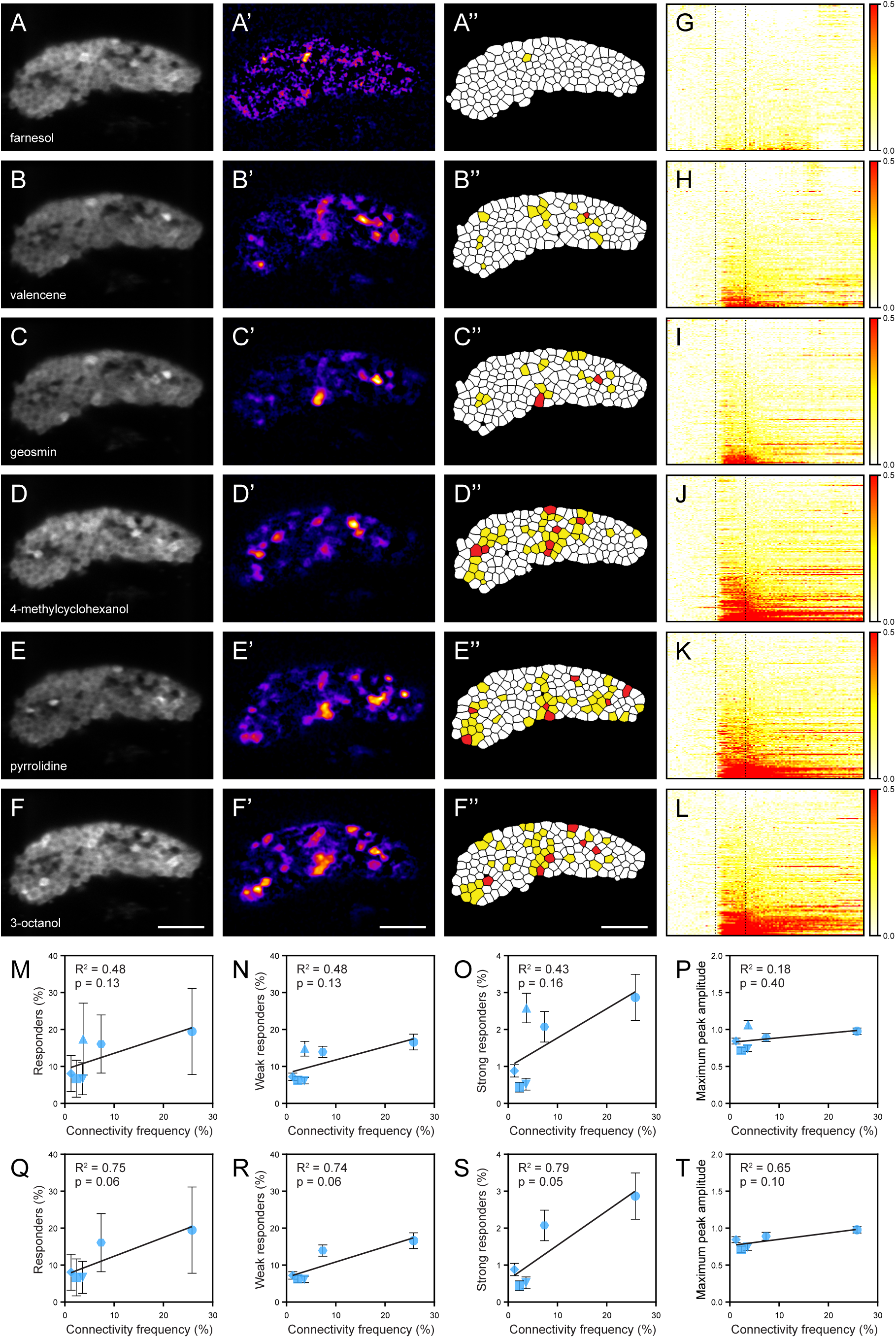
Odor-evoked Kenyon cell responses correspond to connectivity frequency. (A–R) Kenyon cell responses to stimulation with odors activating different numbers of glomeruli. For each odor, panels show: a representative fluorescence image of Kenyon cells expressing GCaMP6f in a single optical plane (A–F), a heat map of ΔF responses following odor presentation (A’-F’), and segmented activation map in which non-responsive Kenyon cells are filled in white, weak responders in yellow, and strong responders in red (A″-F″). Scale bar: 20 μm. (G–L) Representative raster plots showing calcium responses from all recorded Kenyon cells following presentation of each odor: farnesol (G), valencene (H), geosmin (I), 4-methylcyclohexanol (J), pyrrolidine (K), and 3-octanol (L). Each row represents one cell, each column represents one imaging frame, and color intensity indicates ΔF/F amplitude at that frame (scale: 0 to 0.5 standard deviation). The dotted rectangular outline marks odor onset and offset. (M–T) Quantification of Kenyon cell responses plotted against the cumulative connectivity frequency of the glomeruli activated by each odor (Table 2). Panels M–P include all odors; panels Q–T show the same analyses with pyrrolidine excluded. (M,Q) Percentage of all responders. (N,R) Percentage of weak responders. (O,S) Percentage of strong responders. (P,T) Mean maximum peak ΔF/F amplitude. Each data point represents an odor; error bars show standard error of the mean; linear regression lines are shown with R² and p values. Symbols indicate odor identity: square (farnesol), downward triangle (valencene), lozenge (geosmin), hexagon (4-methylcyclohexanol), upward triangle (pyrrolidine), circle (3-octanol). See Table S3 and Figures S14–S17.

Multi-glomerular odors generally evoked broader Kenyon cell responses, with approximately 20% of Kenyon cells responding, whereas most mono-glomerular odors evoked sparser responses, with fewer than 10% responding (Figure 4M). The frequency of responders tended to increase with the cumulative connectivity frequency of the glomeruli activated by an odor, although this correlation did not reach statistical significance (Figures 4M, S15A). As in the optogenetic dataset, the distribution of amplitudes among responders was unimodal with a pronounced positive skew for multi-glomerular odors (Figure S3G-L), and strong responders were more frequent (Table S3, Figure 4O, Figure S15C, Figure S16, Figure S17). By contrast, mean maximum peak amplitude showed no correlation with connectivity frequency consistent with the results we obtained using optogenetic stimulation (Figure 4P, Figure S15D).

Pyrrolidine was a notable exception. This odor activates only two glomeruli, both weakly connected, one of which is the underrepresented VL1 glomerulus. Consistent with our optogenetic results, pyrrolidine activates more Kenyon cells than expected based on its connectivity frequency: it elicited responses in 17.4% ± 9.7 of the Kenyon cells despite activating projection neurons with relatively low connectivity (3.73%), a response magnitude to that evoked by multi-glomerular odors that activate projection neurons with higher connectivity frequencies (Table 2, Figure 4M, Table S3). These broad responses mirrored those evoked by optogenetic stimulation of VL1 projection neurons. When pyrrolidine was excluded from the analysis, the correlations between connectivity frequency and number of responders were stronger (Figure 4Q-T, Figure S15E-H). The VL1 channel therefore departs from the relationship observed for the other odors tested, possibly through the indirect disinhibitory mechanisms considered in the previous section.

Together, these results indicate that odor-evoked Kenyon cell responses correspond to connectivity frequency, except for the VL1 channel.

### Odor-evoked Kenyon cell responses are associated with better learning performance, with a notable exception

Having established that different odors evoked Kenyon cell responses that vary in the number of cells activated, we next asked whether the number of responding Kenyon cells was associated with learning performance. We hypothesized that odors activating more Kenyon cell activation would support stronger memory formation and therefore better learning performance than those activating fewer. To test this, we used classical conditioning paradigms in which flies learned to associate odors with appetitive or aversive stimuli (Krashes and Waddell, 2011a; Krashes and Waddell, 2011b). Learning performance was quantified as a preference index and assessed at both 30 minutes (short-term memory) and 24 hours (long-term memory) after training. We tested the six odors for which Kenyon cell activity had been characterized above.

Consistent with our hypothesis, odors that activated more Kenyon cells generally supported higher learning indices (Figure 5A-D). The multi-glomerular odors 3-octanol and 4-methylcyclohexanol, which activated the largest number of Kenyon cells, produced the highest preference indices across all paradigms, independent of reward type or memory duration. Learning performance correlated positively with both the number of responding Kenyon cells and the cumulative connectivity frequency of activated glomeruli across paradigms (Figure 5E-H,M-P).

**Figure 5.**
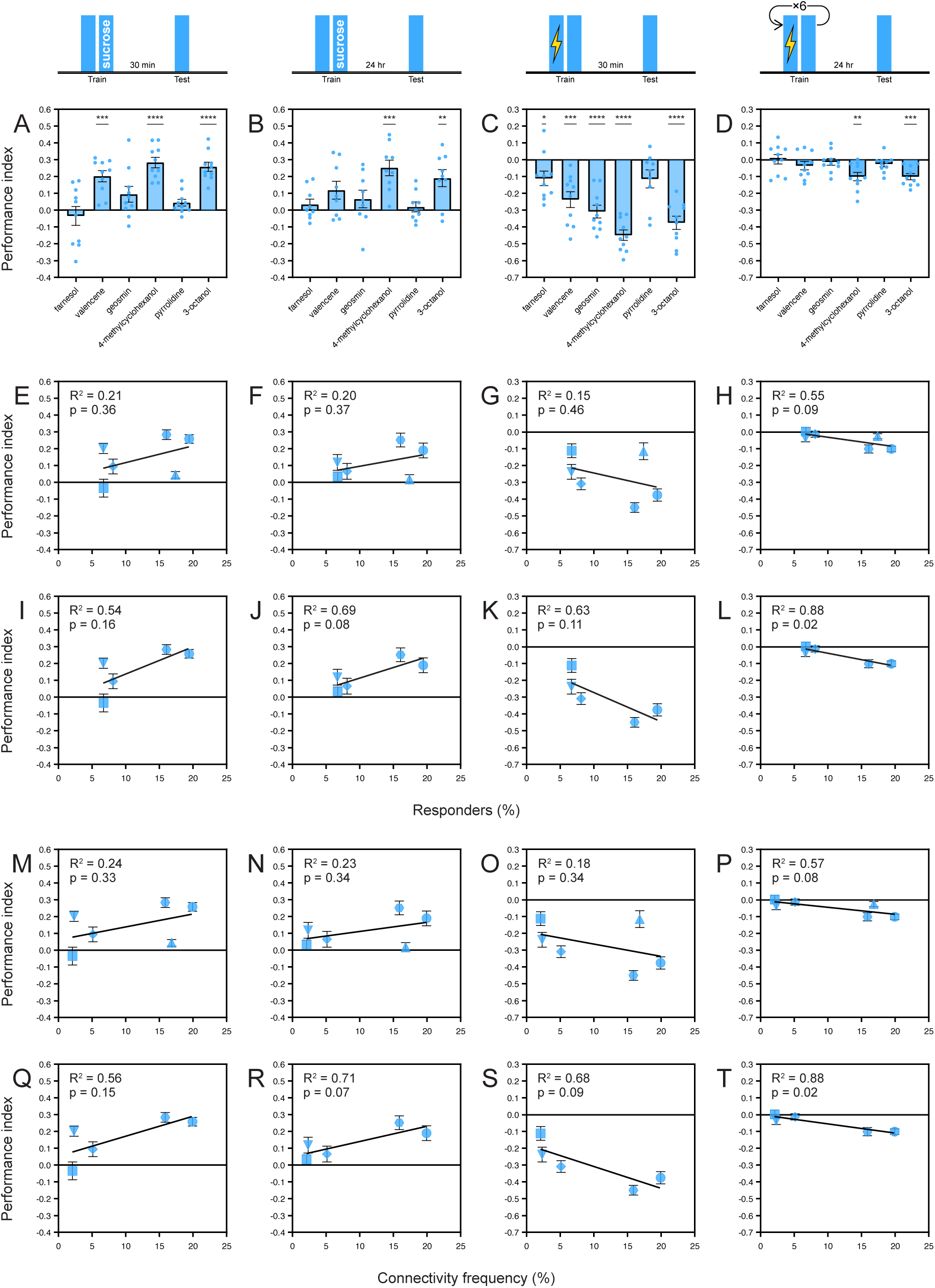
Learning performance correlates with the number of responding Kenyon cells. (A–D) Associative learning performance for six odors measured across four classical conditioning paradigms varying in reinforcement type and memory duration. (A) Appetitive short-term memory assessed 30 minutes after training with sucrose reward. (B) Appetitive long-term memory assessed 24 hours after training. (C) Aversive short-term memory assessed 30 minutes after training with electric shock punishment. (D) Aversive long-term memory assessed 24 hours after training. Each data point represents the mean performance index; error bars show standard error of the mean. Symbols indicate odor identity: square (farnesol), downward triangle (valencene), lozenge (geosmin), hexagon (4-methylcyclohexanol), upward triangle (pyrrolidine), circle (3-octanol). (E–L) Correlation between learning performance and Kenyon cell responses and the number of responding Kenyon cells. The performance index from each paradigm (A–D) is plotted against the percentage of responders for each odor (from Figure 4). Panels E–H include all odors; panels I–L show the same analyses with pyrrolidine excluded. Each data point represents an odor; error bars show standard error of the mean; linear regression lines are shown with R² and p values. Symbol as in (A–D). (M–T) Correlation between learning performance and the cumulative connectivity frequency of the glomeruli activated by each odor (Table 2). Panels M-P include all odors; panels Q–T show the same analyses with pyrrolidine excluded. Each data point represents an odor; error bars show standard error of the mean; linear regression lines are shown with R² and p values. Symbol as in (A–D). See Figure S14–S17.

Pyrrolidine was an exception. Despite activating many Kenyon cells, most likely through VL1 projection neurons, pyrrolidine failed to support learning in any conditioning paradigm (Figure 4E, Table S3, Figure S16, Figure S17). When pyrrolidine was excluded from the analysis, the correlations between learning performance and both the number of responding Kenyon cells and connectivity frequency were stronger (Figure 5I-L,Q-T, Figure S18). The VL1 channel therefore departs from the relationship observed for the other odors tested: the number of responding Kenyon cells and connectivity frequency correspond to learning performance for most odors, but not for pyrrolidine.

Together, these findings are consistent with a relationship between connectivity, function and learning in the mushroom body, in which connectivity biases favor the representation and learning of certain odors, alongside at least one channel that operates outside this relationship.

## DISCUSSION

In the *Drosophila melanogaster* mushroom body, connectivity biases correspond to differences in odor-evoked activity and learning. Projection neurons that form more presynaptic boutons connect more frequently to Kenyon cells and evoke responses in more of them, and the odors they carry tend to support better learning; projection neurons with fewer boutons connect less frequently, evoke sparser responses, and support weaker learning. For most odors, learning performance therefore corresponds to connectivity frequency and to the number of responding Kenyon cells. The VL1 projection neurons are an exception: they activate many Kenyon cells despite low connectivity, yet fail to support learning, indicating that additional circuit mechanisms can operate outside this relationship. Such a relationship between connectivity, function and learning may be one way in which a learning and memory center resolves the trade-off between coding capacity and biological selectivity introduced above.

### VL1 glomerular channel as an exception to the connectivity–function–learning rule

The VL1 glomerular channel is a notable exception to the relationship between connectivity, function and learning. Despite being among the least connected projection neuron types, optogenetic stimulation of VL1 projection neurons produced in unusually large Kenyon cell responses. Pyrrolidine, an odor detected by the VL1 glomerulus, evoked similarly broad responses yet failed to support associative learning in either appetitive or aversive paradigms. The VL1 channel therefore decouples Kenyon cell activity from learning, indicating that additional circuit mechanisms can operate outside this relationship.

At least two non-exclusive mechanisms could account for this. First, the VL1 glomerulus is associated with both excitatory and inhibitory projection neurons, much of whose inhibitory output is directed to the lateral horn rather than the mushroom body (Bates et al., 2020a); whether and how this circuitry contributes to the broad Kenyon cell responses evoked by VL1 stimulation is not known. Second, gating could occur at the level of plasticity rather than activation: VL1 projection neurons may recruit modulatory circuits that block synaptic modification between Kenyon cells and mushroom body output neurons, which would explain the learning failure but not the large number of Kenyon cells. Distinguishing these possibilities will require targeted circuit dissection.

Why the VL1 glomerular channel appears specialized is also unclear. The VL1 glomerulus receives input from Ir75d-expressing olfactory sensory neurons, whose only known ligand is pyrrolidine, a plant secondary metabolite produced as a defensive compound against herbivores (Hartmann and Ober, 2008; Silbering et al., 2011; Prieto-Godino et al., 2017). For odor cues that signal potential toxicity, innate responses may be more adaptive than learned ones, and a circuit that supports robust detection while blocking plasticity downstream would allow rapid responses to such cues without committing them to memory (Stensmyr et al., 2012). Whether and how *D. melanogaster* uses pyrrolidine in its natural environment, and how the VL1 glomerular channel contributes to such behavior, remain open questions.

### Biased connectivity as a common circuit motif in expansion layers

Expansion layers are a recurrent circuit motif in learning and memory centers that can maximize coding capacity by transforming dense sensory inputs into sparse, high-dimensional representations through random connections (Babadi & Sompolinsky, 2014; Litwin-Kumar et al., 2017). Yet in the *D. melanogaster* mushroom body, we and others found that the mushroom body architecture deviates from purely random connectivity, and that biases emphasize certain glomerular channels over others, consistently across electron microscopy connectomes and light-microscopy tracing datasets. Biased connectivity is not restricted to the olfactory modality: visual input to Kenyon cells is also structured rather than uniformly random (Ganguly et al., 2024). These biases may resolve a fundamental trade-off faced by learning and memory centers: maintaining flexibility to represent a wide range of inputs while prioritizing those of greatest behavioral relevance. This architectural feature prevents meaningful information from being diluted among countless equiprobable combinations, allowing the mushroom body architecture to remain both general and selective. In this view, randomness and bias act as complementary architectural features: randomness maximizing coding capacity, and bias allocating that capacity toward ecologically important stimuli.

Hints of biased connectivity as an architectural feature appear in vertebrate expansion layers. In the cerebellum, granule cells receive mossy-fiber inputs that are far from uniform. Mossy fibers originating from related sensorimotor pathways cluster into discrete microzones, and specific body or movement representations occupy disproportionately large territories within the granular layer (Apps & Hawkes, 2009; Nguyen et al., 2023). This arrangement may bias cerebellar learning toward frequently executed or ethologically critical motor patterns, favoring precision and efficiency over uniform sampling. Likewise, in the hippocampal dentate gyrus, inputs from the entorhinal cortex are unevenly distributed: medial entorhinal inputs conveying spatial information and lateral entorhinal inputs carrying object or contextual signals form distinct and form distinct projections (Hargreaves et al., 2005; Amaral et al., 2007; Knierim et al., 2014). Such asymmetries may predispose learning toward spatial or contextual associations, which are essential for survival in freely moving animals. Although the functional consequences of these biases remain to be tested directly, their anatomical prominence suggests that biased connectivity could be a general feature of expansion layers, tuning representation space toward high-value information while retaining capacity for novelty.

Biased connectivity may therefore be a general strategy for resolving the trade-off between coding capacity and selectivity, embedding structured priorities within otherwise random wiring so that learning systems retain both generalization and sensitivity to salient cues. How this balance is tuned across taxa is an open question.

### Interpreting the connectivity, function and learning relationship

Several features of the experimental design temper these conclusions. Neither the optogenetic nor the odor stimuli were controlled for total input drive: Chrimson-mediated activation is sustained and lacks the short-term depression of natural projection neuron output, and odors were not matched for the total olfactory input they evoke, so an odor that recruits more olfactory sensory neurons overall could drive broader responses and better learning for reasons unrelated to connectivity biases. The panel was also small and coarse, as the *split-GAL4* lines cover few projection neuron types, several label pairs or trios of glomeruli, and most activate fewer projection neurons than the coincident input a Kenyon cell typically requires to spike, while the six odors span one to ten glomeruli and provide few points for the correlations reported, several of which did not reach conventional significance. Finite sampling from a circuit that is randomly wired at the single-cell level further inflates the variance of population measures such as the fraction of responding Kenyon cells, which can obscure real correlations at this sample size. The relationships described should therefore be read as tendencies rather than established rules. Glomerular activation was also predicted from a published database rather than measured directly, so the number of glomeruli assigned to each odor is an approximation that does not account for concentration, delivery, or lateral inhibition in the antennal lobe (Olsen and Wilson, 2008; Wilson, 2013). Concentration-matched comparisons, direct measurement of projection neuron drive, and a broader odor panel would be needed to separate glomerular identity from input strength and to test the generality of these relationships.

### Balancing capacity and selectivity

The connectivity biases in the mushroom body, and their functional and behavioral implications, likely reflect evolutionary tuning to the ecological niche specific to *D. melanogaster*. Glomerular channels detecting food sources and mates are among the most overrepresented (Figure 1; Caron et al., 2013), suggesting that biases emphasize stimuli predictive of survival. Expansion layers may therefore not be a mere *tabula rasa* coding device, but a circuit architecture shaped by ecological pressures. No brain, operating with a limited number of neurons and synapses, can encode everything; every brain must prioritize the cues most relevant to survival.

What advantages make this selectivity worth its cost in coding capacity remains unclear. By compressing representational space around biologically relevant dimensions, such biases may support higher-order functions such as stability, generalization, and rapid inference (Zavitz et al., 2021): a biased expansion layer, for instance, may maintain stable and reliable representations even as sensory inputs fluctuate. Additionally, because biases preserve relationships among survival-related stimuli, they may promote generalization across them. Likewise, a circuit already weighted toward biologically meaningful cues could enable faster and more efficient learning. Future behavioral investigations using the experimental framework established here will be essential to determine whether the selectivity afforded by biased connectivity, despite the loss of capacity, supports such higher-order functions. Understanding how evolution tunes this trade-off — how much randomness a brain can afford to lose — will be key to uncovering the architectural features that make learning and memory centers both adaptive and economical.

## MATERIALS AND METHODS

### Fly stocks and husbandry

Flies were reared on standard cornmeal agar medium under standard conditions (25°C, 60% humidity) in incubators that maintain a 12 h light/12h dark cycle (Percival Scientific Inc, Cat#DR36VL). The stocks used and their sources were as follows: *D. melanogaster*: w^1118^;[R13F02-LexA]^attP40^/CyO ; (Bloomington Stock Center, 52460), w^1118^;[13XLexAop2-IVS-GCaMP6f-p10^}su(Hw)attP5^ (Bloomington Stock Center, 44277), w^1118^;[10xUAS-IVS-myrGFP]^Su(Hw)attp5^ (Bloomington Stock Center, 32199), w;;[UAS-Chrimson::mVenus]^attP2^ (D. Anderson, Caltech University), w^1118^;[R54A11-p65.AD]^attP40^; [VT033006-GAL4.DBD]^attP2^ (Bloomington Stock Center, 88835), w;[R44A02-p65ADZp]^attP40^; [VT033006-ZpGdbd]^attP2^ (Janelia Research Campus, SS01165), w;[VT033006-p65ADZp]^attP40^; [R26B04-ZpGdbd]^attP2^ (Janelia Research Campus, SS01306), w;[R65G01-p65ADZp]^attP40^; [R82D04-ZpGdbd]^attP2^ (Janelia Research Campus, MB546B), w;[VT033006-p65ADZp]^attP40^; [R33H11-ZpGdbd]^attP2^ (Janelia Research Campus, SS01318), w;[VT033006-p65ADZp]^attP40^; [R19G08-ZpGdbd]^attP2^ (Janelia Research Campus, SS01265);.

### Connectome analysis

Hemibrain analyses were conducted on the hemibrain:v1.2.1 dataset using a custom query to identify all connections with a weight value of five or greater between the uniglomerular olfactory projection neurons reported in this study and their downstream Kenyon cell synaptic partners. Flywire analyses were performed on the FAFB v783 dataset using a custom script to process the connections_princeton_no_threshold. As in the hemibrain analysis, uniglomerular olfactory projection neurons with at least five synapses onto Kenyon cells were included.

### Quantifying morphological features of projection neurons

Fly brains were dissected at room temperature in phosphate-buffered saline (PBS; Sigma-Aldrich, P5493) and fixed in 2% paraformaldehyde (Electron Microscopy Sciences, 15710) for 45 minutes at room temperature. Following fixation, brains were washed five times in PBST (PBS containing 0.1% Triton X-100; Sigma-Aldrich, T8787) at room temperature and blocked with 5% normal goat serum (Jackson ImmunoResearch Laboratories, AB_2336990) in PBST for 30 minutes at room temperature. For primary antibody incubation, brains were incubated overnight at 4°C in a solution containing mouse anti-nc82 (1:20; Developmental Studies Hybridoma Bank, AB_2314866) and rabbit anti-GFP (1:25; Thermo Fisher Scientific, AB_221569) diluted in 5% normal goat serum/PBST. The following day, brains were washed four times in PBST and incubated overnight at 4°C with secondary antibodies: goat anti-mouse Alexa Fluor 488 (1:500; Thermo Fisher Scientific, AB_2534069) and goat anti-rabbit Alexa Fluor 546 (1:500; Thermo Fisher Scientific, AB_2534077) in 5% normal goat serum/PBST. After secondary antibody incubation, brains were washed four times in PBST and mounted on glass slides (Fisher Scientific, 12-550-143) using VECTASHIELD mounting medium (Vector Laboratories, H-1000). Immunostained preparations were imaged using an LSM 880 confocal microscope (Zeiss).

### Confocal imaging

All confocal images were acquired using a Zeiss LSM 880 microscope equipped with Airyscan technology. Whole-brain images were collected using a Plan-Apochromat 20×/0.8 M27 objective lens with a voxel size of 0.69 × 0.69 × 0.84 μm. Antennal lobe regions were imaged using a Plan-Apochromat 40×/1.3 Oil M27 objective lens with a voxel size of 0.21 × 0.21 × 0.84 μm. High-resolution imaging of mushroom body calyx structures was performed using a Plan-Apochromat 63×/1.4 Oil DIC M27 objective lens with a voxel size of 0.13 × 0.13 × 1.0 μm.

### Cell and bouton quantification

All quantitative analyses of projection neurons were performed by researchers blinded to genotype. For neuronal cell counts, confocal z-stacks of the antennal lobe were analyzed using ImageJ/FIJI software (Schindelin et al., 2012). Individual cells were manually counted by stepping through each optical section of the z-stack and tallying distinct cell bodies using the ‘Cell Counter’ plugin. Mushroom body calyx bouton counts were obtained using the same manual counting approach on confocal z-stacks, with individual boutons tallied using the ‘Cell Counter’ plugin in ImageJ/FIJI. Total bouton volume measurements were performed using FluoRender software (version 2.30.0; Wan et al., 2017). Boutons were selected and segmented using the ‘Paint Brush’ function with ‘Edge Detect’ enabled and the ‘Edge STR’ parameter set to 0.512. Threshold values were adjusted individually for each sample to account for variations in background fluorescence intensity.

### Functional imaging using optogenetic stimulation

Optogenetic stimulation experiments followed the same preparation protocol as odor stimulation experiments, with flies aged two to five days old mounted and imaged using identical microscopy parameters. Prior to experimentation, flies were maintained on fresh food supplemented with 10 μM all-trans-retinal (Sigma-Aldrich, R2500) and protected from light for approximately 24 hours.

Optogenetic activation was delivered using 637 nm light pulses (Coherent, 1196625) configured as five repetitions of five 40 μm spirals targeting the mushroom body calyx over a 100 milliseconds duration. Stimulation protocols consisted of 2.5 seconds baseline recording, followed by optogenetic activation, then 7.5 seconds post-stimulus imaging. This sequence was performed twice at each of the seven planes spanning the mushroom body (6 μm separation), with 1-minute intervals between trials and 1-minute intervals between planes. Images were collected at 496 × 342-pixel resolution (pixel size: 0.19 × 0.19 μm) using a resonant galvanometer scanner at 45.2 fps.

### Functional imaging using odor stimulation

All functional imaging experiments were performed on two-to-five-day-old female flies. Flies were anesthetized on ice for about ten minutes and mounted on a custom platform using tape (Shurtape Technologies, DUC280068). The platform contained an opening connected to a saline reservoir. To minimize movement artifacts, contact points between the head and tape, as well as the legs, were secured using UV-activated resin (Bondic, SK8024). A small window overlying the mushroom body was created in the cuticle, and the exposed brain was continuously immersed in saline containing: 108 mM NaCl, 5 mM KCl, 5 mM HEPES, 5 mM trehalose, 10 mM sucrose, 1 mM NaH₂PO₄, 4 mM NaHCO₃, 2 mM CaCl₂, 4 mM MgCl₂, and 1.7 mM NaOH (pH 7.3).

Odor stimuli — farnesol (Sigma-Aldrich, 43348), (+)-valencene (Sigma-Aldrich, 75056), geosmin (Sigma-Aldrich, UC18), pyrrolidine (Sigma-Aldrich, 69581), 3-octanol (Sigma-Aldrich, 218405), and 4-methylcyclohexanol (Sigma-Aldrich, 153095) — were each diluted to 10^−3^ (v/v) in mineral oil (Sigma-Aldrich, M5904). Filtered carrier air (1.6 l/min) was mixed with odors (0.4 l/min) in a flow-controlled manner using a stimulus controller (Ockenfels Syntech, CS-55). All odors were presented at a single dilution (10⁻³, v/v) and were not matched for the total olfactory input they evoke. The glomeruli activated by each odor were predicted from the DoOR 2.0 database (Münch and Galizia, 2016) rather than measured in this preparation; these predictions do not account for odor concentration or for lateral inhibition in the antennal lobe.

Calcium imaging was performed using an Investigator two-photon laser scanning microscope (Bruker Corporation, RRID:SCR_019807) equipped with a Chameleon Ti:Sapphire laser (Coherent) tuned to 925 nm. Laser power was modulated by Pockels cells (Conoptics, 350-80LA/BK-02) and maintained between 14-24 mW across experiments. Emitted fluorescence was detected using a GaAsP photomultiplier tube (Hamamatsu Photonics). Images were acquired using a Bruker piezo device, imaging across seven focal planes spanning the mushroom body, with planes separated by 6 μm each.

Odors were presented in pseudo-random order using the following protocol: 5 seconds baseline, 3 seconds stimulus presentation, 12 seconds recovery. This sequence was repeated four times per odor with 1-minute inter-trial intervals and 2-minute inter-odor intervals. All six odors were presented to each preparation. Images were collected at 496 × 342-pixel resolution (pixel size: 0.19 × 0.19 μm) using a resonant galvanometer scanner at an effective frame rate of 5 fps per plane.

### Functional imaging processing

Image sequences were motion-corrected using CaImAn-MATLAB (Giovannucci et al. 2019), with parameters adjusted individually based on movement severity. Motion-corrected sequences were maximum intensity projected using ImageJ/FIJI and automatically segmented using a custom-trained cyto3 model in Cellpose (Stringer et al., 2021). Cellpose parameters were: diameter = 18.44, flow threshold = 0, cell probability threshold = −5, and minimum size = 100 pixels. All automated segmentations were manually reviewed, with regions of interest added or removed in cases of obvious segmentation errors. Preparations with poor segmentation performance due to unresolvable motion artifacts were excluded from analysis. Fluorescence changes were quantified as ΔF/F using a custom Python script, where: ΔF/F = (F(t) - F₀) / F₀; F(t) represents raw fluorescence at time point t, and F₀ represents the mean baseline fluorescence calculated from pre-stimulus frames. ΔF/F values were calculated for each trial and concatenated to generate final traces for downstream analysis.

### Appetitive and aversive learning paradigm

Appetitive and aversive learning paradigms were conducted following previously published protocols (Ellis et al., 2024). In short, groups of 60 to 100 male and female flies were collected several hours before experimentation. For aversive learning, flies were maintained on standard food throughout the experiment. For appetitive learning, flies were transferred to vials containing 1% agar 16 to 20 hours before training and kept on agar for the duration of the paradigm. All training was performed in a T-maze apparatus (CelExplorer Labs, TMK-501) with a constant airflow maintained of 0.7 L/min regulated by a flowmeter (Dwyer Instruments, 116011-01).

For aversive conditioning, flies were first exposed to a conditioned stimulus (CS+) paired with electric shocks (12 pulses, 90 V, 0.2 Hz) delivered by a stimulator (Grass Instruments, S48) for one minute, followed by a 45-second rest in ambient air. They were then exposed to the second odor (CS−) for one minute without shock, followed by another 45-second rest period. For appetitive conditioning, flies were first exposed to the CS− in a tube containing dry paper for two minutes, followed by a 30-second rest in ambient air. They were then exposed to the CS+ in a tube lined with paper coated in dried sucrose, followed by another 30-second rest period.

Conditioned stimuli consisted of odors diluted to 10⁻³ (v/v) in mineral oil or mineral oil alone (Sigma-Aldrich, M5904). The odors were farnesol (10⁻³, v/v; Sigma-Aldrich, 43348), 3-octanol (10⁻³, v/v; Sigma-Aldrich, 218405), 4-methylcyclohexanol (10⁻³, v/v; Sigma-Aldrich, 153095), (+)-valencene (10⁻³, v/v; Sigma-Aldrich, 75056), pyrrolidine (10⁻³, v/v; Sigma-Aldrich, 69581), and (±)-geosmin (10⁻³, v/v; Sigma-Aldrich, FR162423).

Following training, flies were transferred to vials containing standard food (aversive paradigm) or 1% agar (appetitive paradigm) and held for either 30 minutes (short-term memory) or 24 hours (long-term memory). During testing, flies were given a choice between T-maze arms containing the CS+ or CS−. A performance index (PI) was calculated as: PI = (CS+ - CS-)/(CS+ + CS-), where CS+ and CS− represent the number of flies in each arm. Each data point represents the average of reciprocal experiments in which CS+ and CS− assignments were switched.

Innate odor preference was measured by allowing flies to choose between tubes containing odor or mineral oil for two minutes. Shock avoidance was assessed by allowing flies to choose between chambers with electrified copper grids (90 V pulses every five seconds for one minute) and non-electrified grids. Sugar preference was measured by offering flies a choice between chambers lined with dry paper or paper coated with dried sucrose.

### Statistical analyses

Each sample corresponds to one preparation from one fly. Sample sizes for each experiment are reported in Tables S2 and S3. For morphological quantification, one hemisphere was analyzed *per* brain, and each brain was treated as an independent sample. Error bars in all figures represent the standard deviation, and individual data points are shown.

To determine whether glomeruli were connected to Kenyon cells at frequencies that deviate from those expected under random connectivity, glomerular representation was tested against a null distribution. For the large-scale tracing datasets, a shuffle-based approach was used: for each connectivity matrix, 1,000 uniform shuffle matrices were generated in which the total number of projection neuron–Kenyon cell connections was preserved but connections were randomly reassigned, and 1,000 biased shuffle matrices were generated in which connections were reassigned while preserving the measured connectivity frequency of each glomerulus and the distribution of input counts per Kenyon cell, as described previously (Caron et al., 2013; Hayashi et al., 2022; Ellis et al., 2024). A glomerulus was classified as over- or underrepresented if its connectivity frequency differed significantly from the value expected under uniform connectivity (Fisher’s exact test, with a Benjamini-Hochberg false-discovery-rate correction of 10%). For the electron microscopy–based connectomes, each glomerulus was tested against the frequency expected under uniform connectivity using a two-sided binomial test (p < 0.05).

To quantify Kenyon cell responses to optogenetic and odor stimulation, Kenyon cells were classified as responders if the post-stimulation mean ΔF/F exceeded twice the standard deviation of the ΔF/F measured during the pre-stimulation period. Responders were further classified as weak or strong by k-means clustering (k = 2 clusters) applied to the maximum ΔF/F of the trial-averaged trace, using the scikit-learn package (Pedregosa et al., 2011).

To assess whether connectivity frequency relates to Kenyon cell responses and to learning performance, correlations were computed by Pearson correlation with simple linear regression in GraphPad Prism version 10.4.0 (GraphPad Software, San Diego, CA). Given the small number of projection neuron lines and odors, these correlations are exploratory, and effect sizes are reported alongside p-values throughout. To determine whether flies formed associative memories, behavioral performance indices were tested against zero using a one-sample t-test, with significance defined as p < 0.05. Significance levels are reported as follows: p < 0.05 (*), p < 0.01 (**), p < 0.001 (***), p < 0.0001 (****), and ns (non-significant).

## ACKNOWLEDGMENTS

We thank members of the Caron laboratory for their comments on the manuscript and discussions throughout the project. We are grateful to Dr. Florian Maderspacher for thoughtful discussions, and careful feedback. We thank Yoshi Aso for sharing *split-GAL4* lines prior to their publication; Adam Lin and Sylvia Yang for preparing the standard cornmeal agar medium; we also want to acknowledge Adam Weinbrom, Hayley Smihula, Ashley Platt, Miles Jacob, and Lia Beatty for providing lab management during the project. We also thank the Cell Imaging Core at the University of Utah for use of the Zeiss LSM 880 microscope. We acknowledge the use of the following software and databases: Fiji (Schindelin et al. 2012), Cellpose (Stringer et al. 2021), DoOR 2.0 (Münch and Galizia, 2016), CaImAn (Giovannucci et al. 2019), FluoRender (Wan et al., 2017), Matplotlib (Hunter, 2007), Navis (Bates et al., 2020b). We thank the Bloomington Drosophila Stock Center (NIH P40OD018537) for providing all the *Drosophila* stocks.

## AUTHOR CONTRIBUTIONS

A.J.M. and S.J.C.C. conceived the project.

D.B.A. and P.A. performed and analyzed the anatomical experiments under the supervision of A.J.M.

A.J.M. performed and analyzed the functional imaging experiments.

J.M.U., P.A. and A.R.B. performed and analyzed the behavioral experiments.

L.B. and A.J.M. curated and organized all the experimental dataset.

L.B. prepared all the figures under the supervision of A.J.M. and S.J.C.C.

A.J.M. and S.J.C.C. wrote the manuscript with input from all authors.

A.R.B. verified statistical analyses and cross-checked the references and source data during revision.

## Supplemental information

**Table S1.** Connectivity frequencies across datasets. Related to Figure 1. The table reports connectivity frequencies for each glomerulus across electron microscopy connectomes (Scheffer et al., 2020; Dorkenwald et al., 2024) and large-scale tracing datasets (Caron et al., 2013; Hayashi et al., 2022; Ellis et al., 2024). Values are reported as means with standard error of the mean.

| Glomerulus | Scheffer<br>*et al.* 2020 (♀) | Dorkenwald<br>*et al.* 2024 (♀) | Average<br>connectivity | Caron *et al.*<br>2013 (♂) | Hayashi *et al.*<br>2022 (♀) | Ellis *et al.*<br>2024 (♀) | Ellis *et al.*<br>2024 (♂) | Average<br>connectivity |
| --- | --- | --- | --- | --- | --- | --- | --- | --- |
| *Average* |  |  | **1.96** |  |  |  |  | **1.98** |
| D | 1.92 | 1.68 | **1.79** | 1.10 | 2.39 | 2.32 | 1.62 | **1.86** |
| DA1 | 3.09 | 3.27 | **3.18** | 5.50 | 3.79 | 4.06 | 4.58 | **4.48** |
| DA2 | 1.49 | 1.02 | **1.25** | 0.47 | 0.14 | 0.58 | 0.30 | **0.37** |
| DA3 | 0.36 | 0.32 | **0.34** | 0.00 | 0.28 | 0.44 | 0.74 | **0.36** |
| DA4l | 0.85 | 1.28 | **1.07** | 0.94 | 0.56 | 0.87 | 1.77 | **1.04** |
| DA4m | 0.30 | 0.39 | **0.35** | 0.31 | 0.56 | 0.73 | 0.44 | **0.51** |
| DC1 | 3.41 | 3.79 | **3.60** | 5.19 | 5.48 | 3.34 | 3.25 | **4.31** |
| DC2 | 2.41 | 1.72 | **2.06** | 2.67 | 2.53 | 1.89 | 1.77 | **2.22** |
| DC3 | 2.4 | 2.19 | **2.30** | 5.35 | 4.07 | 4.50 | 3.10 | **4.26** |
| DC4 | 0.59 | 0.76 | **0.67** | 0.63 | 1.83 | 2.32 | 2.36 | **1.79** |
| DL1 | 3.44 | 2.80 | **3.12** | 4.72 | 4.49 | 4.35 | 3.55 | **4.28** |
| DL2d | 1.95 | 1.65 | **1.80** | 0.63 | 1.54 | 1.02 | 1.48 | **1.17** |
| DL2v | 1.38 | 2.37 | **1.87** | 0.31 | 1.26 | 1.31 | 1.03 | **0.98** |
| DL3 | 0.41 | 0.52 | **0.47** | 1.89 | 1.54 | 0.44 | 0.89 | **1.19** |
| DL4 | 0.41 | 0.28 | **0.34** | 0.94 | 0.56 | 0.73 | 0.59 | **0.71** |
| DL5 | 1.41 | 1.52 | **1.47** | 3.62 | 2.39 | 3.05 | 3.55 | **3.15** |
| DM1 | 3.81 | 3.83 | **3.82** | 1.73 | 2.25 | 1.45 | 1.33 | **1.69** |
| DM2 | 3.13 | 3.09 | **3.11** | 1.10 | 1.69 | 1.60 | 2.95 | **1.83** |
| DM3 | 2.13 | 2.04 | **2.09** | 0.63 | 0.84 | 1.89 | 1.18 | **1.14** |
| DM4 | 2.93 | 3.00 | **2.96** | 0.63 | 0.98 | 1.74 | 1.18 | **1.13** |
| DM5 | 1.97 | 1.94 | **1.95** | 0.63 | 2.25 | 1.16 | 1.03 | **1.27** |
| DM6 | 3.09 | 2.50 | **2.80** | 3.14 | 2.11 | 3.05 | 3.40 | **2.92** |
| DP1l | 3.19 | 2.75 | **2.97** | 1.42 | 1.69 | 0.44 | 1.18 | **1.18** |
| DP1m | 4.91 | 4.90 | **4.91** | 3.30 | 5.20 | 2.18 | 4.28 | **3.74** |
| V | 1.93 | 1.89 | **1.91** | 0.31 | 0.00 | 0.29 | 0.15 | **0.19** |
| VA1d | 1.37 | 1.85 | **1.61** | 3.77 | 2.67 | 5.08 | 2.81 | **3.58** |
| VA1v | 0.98 | 1.55 | **1.27** | 3.30 | 1.83 | 3.19 | 3.55 | **2.97** |
| VA2 | 2.89 | 2.99 | **2.94** | 1.42 | 2.25 | 2.32 | 0.30 | **1.57** |
| VA3 | 2.35 | 2.58 | **2.46** | 2.36 | 3.23 | 3.19 | 2.36 | **2.79** |
| VA4 | 2.42 | 2.25 | **2.34** | 1.57 | 0.98 | 1.16 | 0.89 | **1.15** |
| VA5 | 1.80 | 1.64 | **1.72** | 2.83 | 2.53 | 2.90 | 2.95 | **2.80** |
| VA6 | 2.29 | 2.45 | **2.37** | 4.09 | 2.67 | 4.06 | 3.40 | **3.55** |
| VA7l | 1.34 | 1.20 | **1.27** | 0.79 | 1.54 | 1.16 | 2.36 | **1.46** |
| VA7m | 2.38 | 2.16 | **2.27** | 1.42 | 2.11 | 1.16 | 1.48 | **1.54** |
| VC1 | 1.29 | 1.16 | **1.23** | 1.57 | 1.40 | 1.60 | 1.77 | **1.59** |
| VC2 | 1.68 | 1.62 | **1.65** | 1.42 | 0.98 | 1.16 | 1.62 | **1.30** |
| VC3 | 3.07 | 3.23 | **3.15** | 4.40 | 2.95 | 2.90 | 4.43 | **3.67** |
| VC4 | 2.20 | 1.68 | **1.94** | 1.42 | 2.39 | 1.74 | 2.07 | **1.90** |
| VC5 | 2.63 | 2.15 | **2.39** | 0.00 | 1.97 | n/a | n/a | **0.98** |
| VL1 | 0.40 | 0.44 | **0.42** | 0.00 | 0.56 | 0.44 | 0.44 | **0.36** |
| VL2a | 2.09 | 1.99 | **2.04** | 3.62 | 1.54 | 4.21 | 3.69 | **3.27** |
| VL2p | 2.75 | 2.21 | **2.48** | 3.14 | 2.11 | 3.05 | 2.07 | **2.59** |
| VM1 | 0.81 | 1.24 | **1.03** | 2.04 | 1.12 | 2.03 | 1.33 | **1.63** |
| VM2 | 1.56 | 1.52 | **1.54** | 0.31 | 1.69 | 0.73 | 0.59 | **0.83** |
| VM3 | 1.65 | 2.01 | **1.83** | 0.63 | 0.98 | 0.58 | 1.48 | **0.92** |
| VM4 | 1.02 | 1.03 | **1.02** | 1.26 | 1.97 | 1.74 | 2.81 | **1.94** |
| VM5d | 3.25 | 3.37 | **3.31** | 4.09 | 2.67 | 2.76 | 2.81 | **3.08** |
| VM5v | 1.36 | 1.81 | **1.58** | 3.62 | 2.39 | 3.63 | 4.73 | **3.59** |
| VM6 | 1.45 | 1.64 | **1.54** | 0.63 | 1.83 | n/a | n/a | **1.23** |
| VM7d | 1.10 | 1.53 | **1.32** | 2.67 | 1.97 | 3.05 | 2.36 | **2.51** |
| VM7v | 0.99 | 1.18 | **1.09** | 0.47 | 1.26 | 0.44 | 0.00 | **0.54** |

**Table S2.** Kenyon cell response frequencies for optogenetic experiments. Related to Figure 3. The table shows, for each *split-GAL4* line, the total number of Kenyon cells recorded per sample, along with the percentages of non-responders, weak responders, and strong responders for individual samples. The first row provides summary statistics for the entire dataset, including the total percentage of responders across all samples. Values are reported as means with standard error of the mean.

| Total Kenyon cells |  | Frequency of non-responders / weak responders / strong responders (%) |  |  |  |  |  |  |  |  |  |  |
| --- | --- | --- | --- | --- | --- | --- | --- | --- | --- | --- | --- | --- |
|  | VL1 |  | DL3 |  | VM4 |  | DA1 |  | DA4I, DC3 |  | VM5v, VM7d, VM7v |  |
| **Average** | **502** | **87.0 / 10.7 / 2.3** | **506** | **96.4 / 3.5 / 0.1** | **449** | **89.5 / 9.8 / 0.7** | **492** | **88.5 / 9.7 / 1.8** | **420** | **84.1 / 14.5 / 1.4** | **385** | **90.4 / 8.4 / 1.2** |
| *Responders (%)* | *13.0 ± 8.4* |  | *3.6 ± 1.6* |  | *10.5 ± 7.1* |  | *11.5 ± 6.8* |  | *15.9 ± 9.1* |  | *9.6 ± 7.6* |  |
| Sample 1 | 432 | 85.9 / 12.7 / 1.4 | 328 | 97.0 / 3.0 / 0.0 | 439 | 83.4 / 16.6 / 0.0 | 558 | 80.5 / 16.5 / 3.0 | 354 | 79.1 / 20.6 / 0.3 | 423 | 87.2 / 11.8 / 0.9 |
| Sample 2 | 570 | 91.6 / 7.2 / 1.2 | 290 | 95.5 / 4.5 / 0.0 | 396 | 90.4 / 8.8 / 0.8 | 699 | 81.8 / 15.2 / 3.0 | 276 | 73.9 / 22.5 / 3.6 | 197 | 94.4 / 5.1 / 0.5 |
| Sample 3 | 432 | 91.4 / 8.1 / 0.5 | 307 | 96.4 / 3.6 / 0.0 | 520 | 95.0 / 4.6 / 0.4 | 187 | 96.3 / 3.2 / 0.5 | 549 | 85.8 / 13.8 / 0.4 | 336 | 99.4 / 0.6 / 0.0 |
| Sample 4 | 552 | 94.9 / 4.3 / 0.7 | 452 | 94.0 / 6.0 / 0.0 | 407 | 97.1 / 2.7 / 0.2 | 579 | 84.3 / 13.6 / 2.1 | 658 | 81.8 / 16.1 / 2.1 | 431 | 87.2 / 12.5 / 0.2 |
| Sample 5 | 500 | 88.6 / 9.8 / 1.6 | 731 | 94.8 / 5.1 / 0.1 | 310 | 93.2 / 6.5 / 0.3 | 611 | 97.1 / 2.5 / 0.5 | 502 | 93.8 / 5.4 / 0.8 | 509 | 86.4 / 11.6 / 2.0 |
| Sample 6 | 500 | 81.6 / 13.8 / 4.6 | 550 | 96.4 / 3.3 / 0.4 | 671 | 90.0 / 8.9 / 1.0 | 409 | 88.5 / 10.8 / 0.7 | 496 | 67.5 / 28.8 / 3.6 | 384 | 77.9 / 17.4 / 4.7 |
| Sample 7 | 753 | 92.7 / 6.5 / 0.8 | 609 | 97.7 / 2.3 / 0.0 | 625 | 75.0 / 23.0 / 1.9 | 480 | 80.8 / 15.2 / 4.0 | 362 | 95.9 / 4.1 / 0.0 | 448 | 94.2 / 3.6 / 2.2 |
| Sample 8 | 385 | 91.2 / 5.5 / 3.4 | 557 | 98.6 / 1.4 / 0.0 | 222 | 91.9 / 7.2 / 0.9 | 435 | 93.3 / 6.4 / 0.2 | 210 | 87.1 / 11.9 / 1.0 | 281 | 94.3 / 5.7 / 0.0 |
| Sample 9 | 416 | 86.3 / 10.6 / 3.1 | 463 | 98.9 / 1.1 / 0.0 |  |  | 474 | 93.9 / 4.2 / 1.9 | 445 | 93.7 / 6.3 / 0.0 | 355 | 76.1 / 21.7 / 2.3 |
| Sample 10 | 475 | 65.7 / 28.6 / 5.7 | 772 | 94.8 / 5.2 / 0.0 |  |  |  |  | 347 | 82.1 / 15.6 / 2.3 | 507 | 93.7 / 5.3 / 1.0 |
| Sample 11 |  |  |  |  |  |  |  |  |  |  | 437 | 95.7 / 3.9 / 0.5 |
| Sample 12 |  |  |  |  |  |  |  |  |  |  | 309 | 98.7 / 1.3 / 0.0 |
| Cumulative Kenyon cells |  | Cumulative weak responders / strong responders |  |  |  |  |  |  |  |  |  |  |
| **Cumulative** | **5015** | **523 / 109** | **5059** | **183 / 3** | **3590** | **383 / 28** | **4432** | **463 / 86** | **4199** | **609 / 59** | **4617** | **399 / 59** |

**Table S3.**
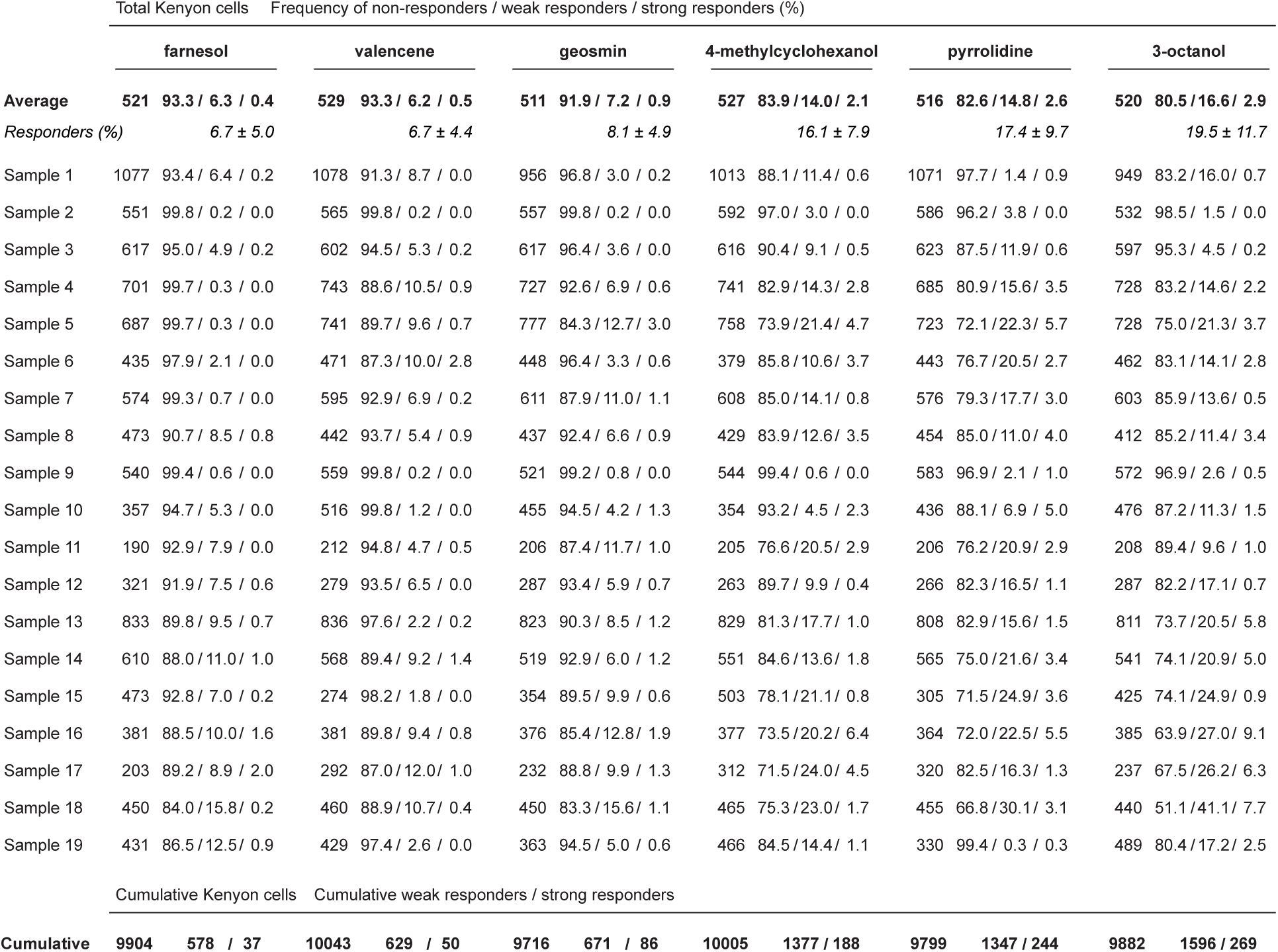
Kenyon cell response frequencies for odor experiments. Related to Figure 4. The table shows, for each odor, the total number of Kenyon cells recorded per sample, along with the percentages of non-responders, weak responders, and strong responders for individual samples. The first row provides summary statistics for the entire dataset, including the total percentage of responders across all samples. Values are reported as means with standard error of the mean.

|  | farnesol |  |  | valencene |  |  | geosmin |  |  | 4-methylcyclohexanol |  |  | pyrrolidine |  |  | 3-octanol |  |
| --- | --- | --- | --- | --- | --- | --- | --- | --- | --- | --- | --- | --- | --- | --- | --- | --- | --- |
| **Average** | **521** | **93.3 / 6.3 / 0.4** |  | **529** | **93.3 / 6.2 / 0.5** |  | **511** | **91.9 / 7.2 / 0.9** |  | **527** | **83.9 / 14.0 / 2.1** |  | **516** | **82.6 / 14.8 / 2.6** |  | **520** | **80.5 / 16.6 / 2.9** |
| *Responders (%)* |  | *6.7 ± 5.0* |  |  | *6.7 ± 4.4* |  |  | *8.1 ± 4.9* |  |  | *16.1 ± 7.9* |  |  | *17.4 ± 9.7* |  |  | *19.5 ± 11.7* |
| Sample 1 | 1077 | 93.4 / 6.4 / 0.2 |  | 1078 | 91.3 / 8.7 / 0.0 |  | 956 | 96.8 / 3.0 / 0.2 |  | 1013 | 88.1 / 11.4 / 0.6 |  | 1071 | 97.7 / 1.4 / 0.9 |  | 949 | 83.2 / 16.0 / 0.7 |
| Sample 2 | 551 | 99.8 / 0.2 / 0.0 |  | 565 | 99.8 / 0.2 / 0.0 |  | 557 | 99.8 / 0.2 / 0.0 |  | 592 | 97.0 / 3.0 / 0.0 |  | 586 | 96.2 / 3.8 / 0.0 |  | 532 | 98.5 / 1.5 / 0.0 |
| Sample 3 | 617 | 95.0 / 4.9 / 0.2 |  | 602 | 94.5 / 5.3 / 0.2 |  | 617 | 96.4 / 3.6 / 0.0 |  | 616 | 90.4 / 9.1 / 0.5 |  | 623 | 87.5 / 11.9 / 0.6 |  | 597 | 95.3 / 4.5 / 0.2 |
| Sample 4 | 701 | 99.7 / 0.3 / 0.0 |  | 743 | 88.6 / 10.5 / 0.9 |  | 727 | 92.6 / 6.9 / 0.6 |  | 741 | 82.9 / 14.3 / 2.8 |  | 685 | 80.9 / 15.6 / 3.5 |  | 728 | 83.2 / 14.6 / 2.2 |
| Sample 5 | 687 | 99.7 / 0.3 / 0.0 |  | 741 | 89.7 / 9.6 / 0.7 |  | 777 | 84.3 / 12.7 / 3.0 |  | 758 | 73.9 / 21.4 / 4.7 |  | 723 | 72.1 / 22.3 / 5.7 |  | 728 | 75.0 / 21.3 / 3.7 |
| Sample 6 | 435 | 97.9 / 2.1 / 0.0 |  | 471 | 87.3 / 10.0 / 2.8 |  | 448 | 96.4 / 3.3 / 0.6 |  | 379 | 85.8 / 10.6 / 3.7 |  | 443 | 76.7 / 20.5 / 2.7 |  | 462 | 83.1 / 14.1 / 2.8 |
| Sample 7 | 574 | 99.3 / 0.7 / 0.0 |  | 595 | 92.9 / 6.9 / 0.2 |  | 611 | 87.9 / 11.0 / 1.1 |  | 608 | 85.0 / 14.1 / 0.8 |  | 576 | 79.3 / 17.7 / 3.0 |  | 603 | 85.9 / 13.6 / 0.5 |
| Sample 8 | 473 | 90.7 / 8.5 / 0.8 |  | 442 | 93.7 / 5.4 / 0.9 |  | 437 | 92.4 / 6.6 / 0.9 |  | 429 | 83.9 / 12.6 / 3.5 |  | 454 | 85.0 / 11.0 / 4.0 |  | 412 | 85.2 / 11.4 / 3.4 |
| Sample 9 | 540 | 99.4 / 0.6 / 0.0 |  | 559 | 99.8 / 0.2 / 0.0 |  | 521 | 99.2 / 0.8 / 0.0 |  | 544 | 99.4 / 0.6 / 0.0 |  | 583 | 96.9 / 2.1 / 1.0 |  | 572 | 96.9 / 2.6 / 0.5 |
| Sample 10 | 357 | 94.7 / 5.3 / 0.0 |  | 516 | 99.8 / 1.2 / 0.0 |  | 455 | 94.5 / 4.2 / 1.3 |  | 354 | 93.2 / 4.5 / 2.3 |  | 436 | 88.1 / 6.9 / 5.0 |  | 476 | 87.2 / 11.3 / 1.5 |
| Sample 11 | 190 | 92.9 / 7.9 / 0.0 |  | 212 | 94.8 / 4.7 / 0.5 |  | 206 | 87.4 / 11.7 / 1.0 |  | 205 | 76.6 / 20.5 / 2.9 |  | 206 | 76.2 / 20.9 / 2.9 |  | 208 | 89.4 / 9.6 / 1.0 |
| Sample 12 | 321 | 91.9 / 7.5 / 0.6 |  | 279 | 93.5 / 6.5 / 0.0 |  | 287 | 93.4 / 5.9 / 0.7 |  | 263 | 89.7 / 9.9 / 0.4 |  | 266 | 82.3 / 16.5 / 1.1 |  | 287 | 82.2 / 17.1 / 0.7 |
| Sample 13 | 833 | 89.8 / 9.5 / 0.7 |  | 836 | 97.6 / 2.2 / 0.2 |  | 823 | 90.3 / 8.5 / 1.2 |  | 829 | 81.3 / 17.7 / 1.0 |  | 808 | 82.9 / 15.6 / 1.5 |  | 811 | 73.7 / 20.5 / 5.8 |
| Sample 14 | 610 | 88.0 / 11.0 / 1.0 |  | 568 | 89.4 / 9.2 / 1.4 |  | 519 | 92.9 / 6.0 / 1.2 |  | 551 | 84.6 / 13.6 / 1.8 |  | 565 | 75.0 / 21.6 / 3.4 |  | 541 | 74.1 / 20.9 / 5.0 |
| Sample 15 | 473 | 92.8 / 7.0 / 0.2 |  | 274 | 98.2 / 1.8 / 0.0 |  | 354 | 89.5 / 9.9 / 0.6 |  | 503 | 78.1 / 21.1 / 0.8 |  | 305 | 71.5 / 24.9 / 3.6 |  | 425 | 74.1 / 24.9 / 0.9 |
| Sample 16 | 381 | 88.5 / 10.0 / 1.6 |  | 381 | 89.8 / 9.4 / 0.8 |  | 376 | 85.4 / 12.8 / 1.9 |  | 377 | 73.5 / 20.2 / 6.4 |  | 364 | 72.0 / 22.5 / 5.5 |  | 385 | 63.9 / 27.0 / 9.1 |
| Sample 17 | 203 | 89.2 / 8.9 / 2.0 |  | 292 | 87.0 / 12.0 / 1.0 |  | 232 | 88.8 / 9.9 / 1.3 |  | 312 | 71.5 / 24.0 / 4.5 |  | 320 | 82.5 / 16.3 / 1.3 |  | 237 | 67.5 / 26.2 / 6.3 |
| Sample 18 | 450 | 84.0 / 15.8 / 0.2 |  | 460 | 88.9 / 10.7 / 0.4 |  | 450 | 83.3 / 15.6 / 1.1 |  | 465 | 75.3 / 23.0 / 1.7 |  | 455 | 66.8 / 30.1 / 3.1 |  | 440 | 51.1 / 41.1 / 7.7 |
| Sample 19 | 431 | 86.5 / 12.5 / 0.9 |  | 429 | 97.4 / 2.6 / 0.0 |  | 363 | 94.5 / 5.0 / 0.6 |  | 466 | 84.5 / 14.4 / 1.1 |  | 330 | 99.4 / 0.3 / 0.3 |  | 489 | 80.4 / 17.2 / 2.5 |
| Cumulative Kenyon cells Cumulative weak responders / strong responders |  |  |  |  |  |  |  |  |  |  |  |  |  |  |  |  |  |
| **Cumulative** | **9904** | **578 / 37** |  | **10043** | **629 / 50** |  | **9716** | **671 / 86** |  | **10005** | **1377 / 188** |  | **9799** | **1347 / 244** |  | **9882** | **1596 / 269** |

**Figure S1.**
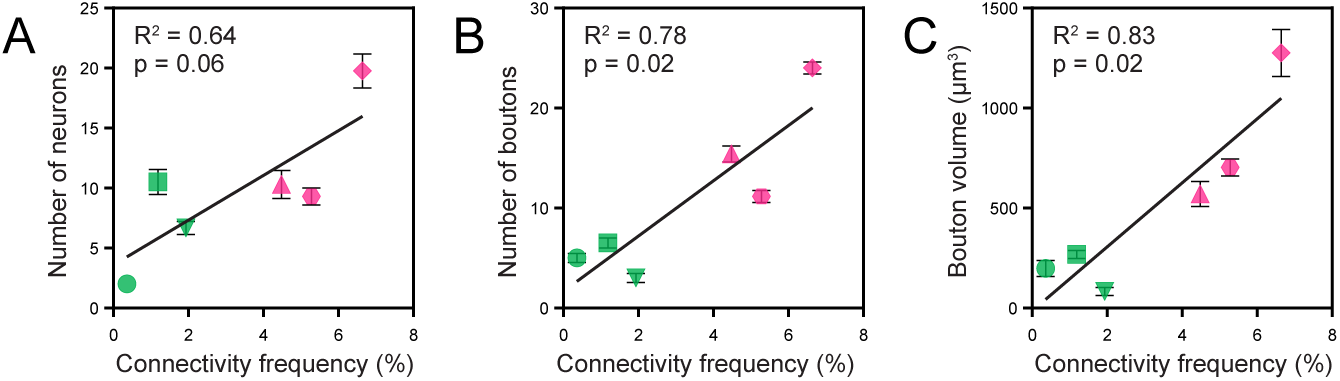
Morphological features correlate with connectivity frequency in large-scale tracing datasets. Related to Figure 2. (A–C) Quantification of morphological features versus average connectivity frequency from large-scale tracing datasets. (A) Number of projection neurons per *split-GAL4 line*. (B) Total bouton number in calyx. (C) Total bouton volume. Each data point represents a *split-GAL4* line; error bars show standard error of the mean; linear regression lines are shown with R² and p values. Symbols indicate projection neuron identity: circle (VL1), square (DL3), downward triangle (VM4), upward triangle (DA1), hexagon (DA4l+DC3), diamond (VM5d+VM7d+VM7v). Colors indicate degree of representation: pink (overrepresented) and green (underrepresented).

**Figure S2.**
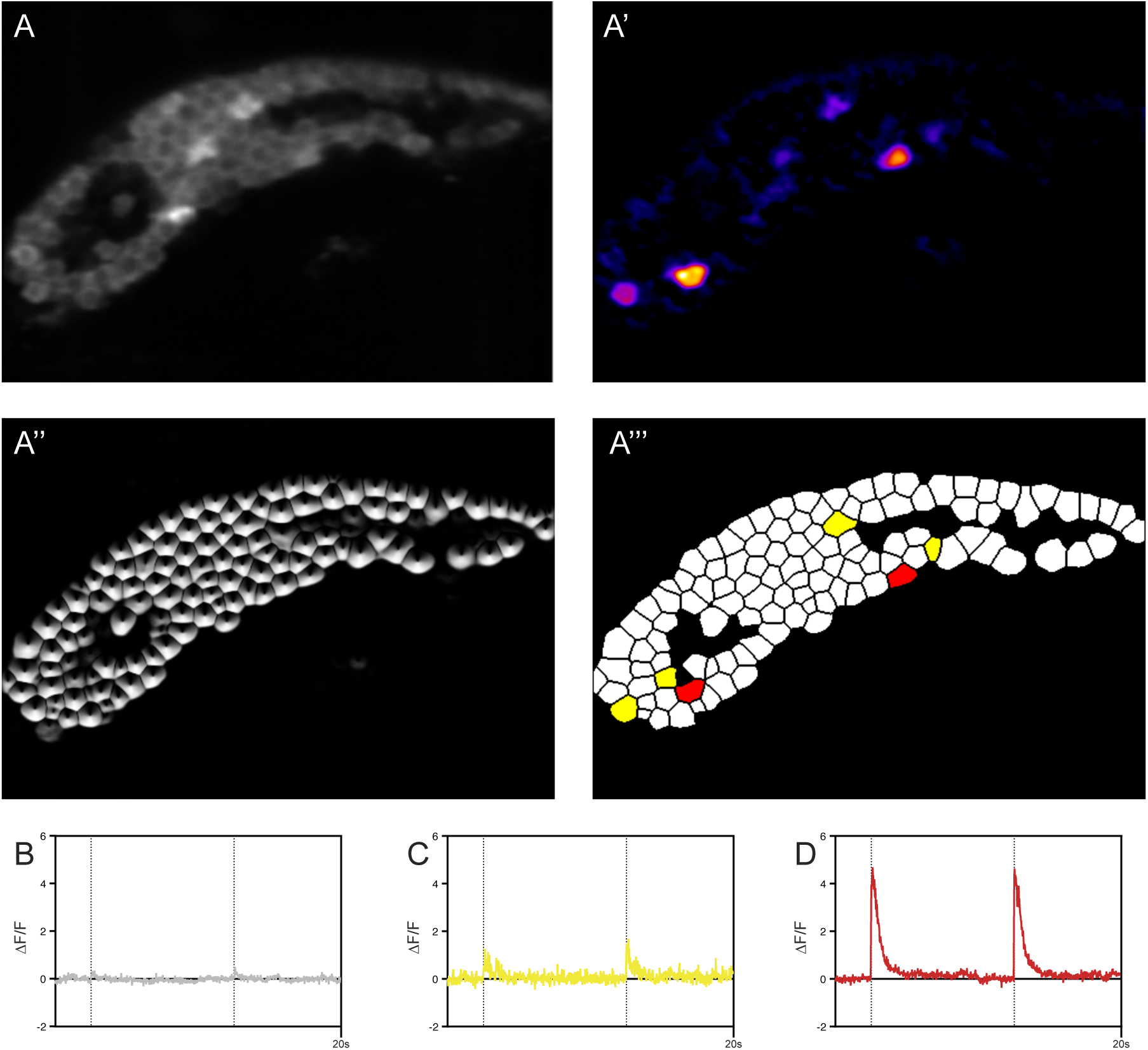
Classification of Kenyon cell responses to optogenetic stimulation. Related to Figure 3. (A) Representative fluorescence image of Kenyon cells expressing GCaMP6f during optogenetic stimulation with *split-GAL4* line targeting the expression of *UAS-Chrimson* in VL1 projection neurons. (A’) Heat map of ΔF responses following optogenetic stimulation. (A’’) Segmented image showing individual Kenyon cell boundaries. (A’’’) Classification map showing non-responders (white), weak responders (yellow), and strong responders (red). (B–D) Representative calcium traces (ΔF/F) from a non-responder (B), weak responder (C), and strong responder (D). Black lines indicate two stimuli presentations *per* sample.

**Figure S3.**
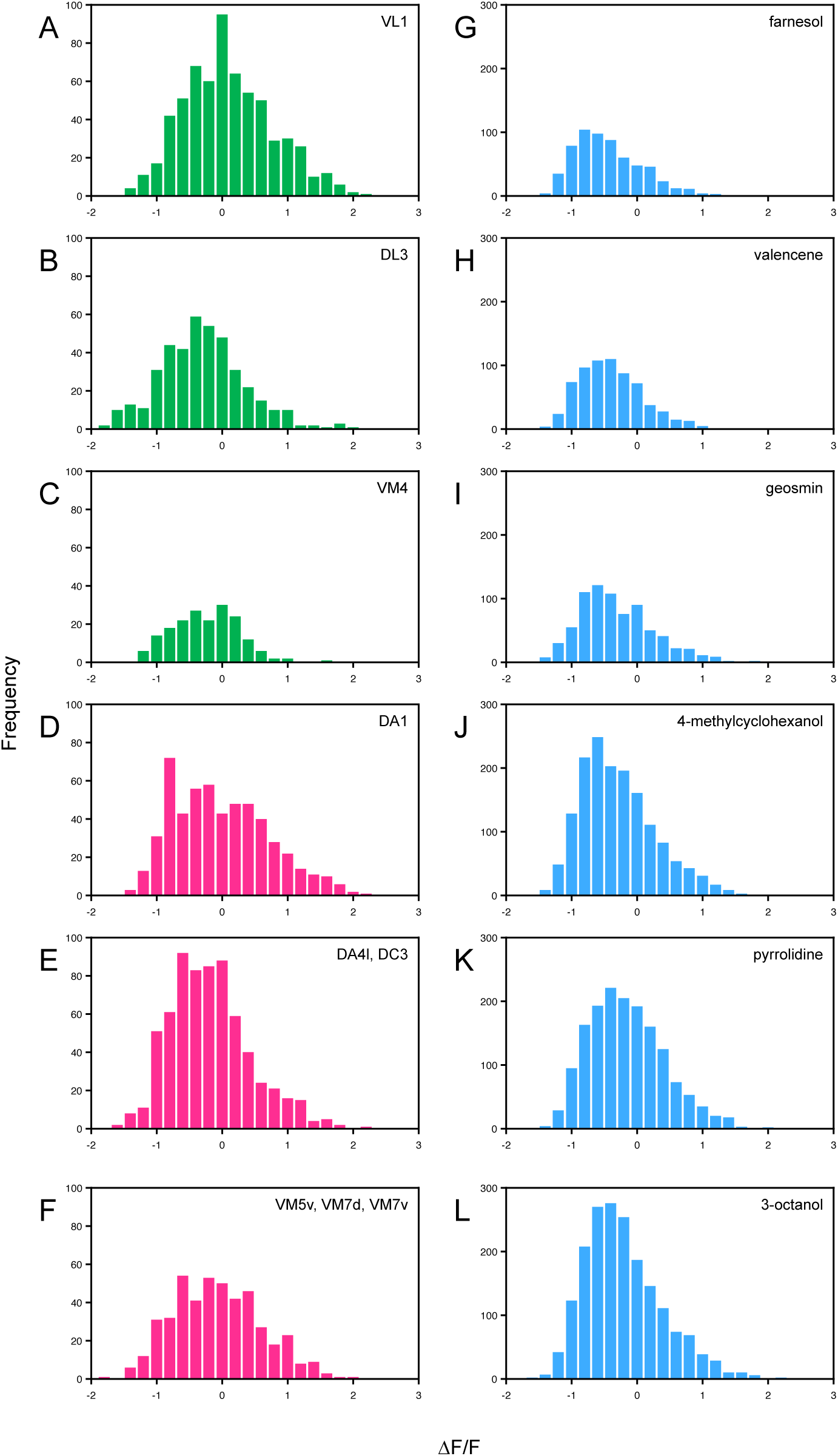
Distribution of response amplitudes across recorded Kenyon cells. Related to Figures 3 and 4 (A–F) Histograms showing the natural logarithm of the maximum ΔF/F amplitude distributions among all Kenyon cell responders following optogenetic stimulation of each *split-GAL4* line: VL1 (A), DL3 (B), VM4 (C), DA1 (D), DA4l+DC3 (E), VM5d+VM7d+VM7v (F). (G–L) Histograms showing the natural logarithm of the maximum ΔF/F amplitude distributions among all Kenyon cell responders following odor presentation: farnesol (G), valencene (H), geosmin (I), 4-methylcyclohexanol (J), pyrrolidine (K), 3-octanol (L).

**Figure S4.**
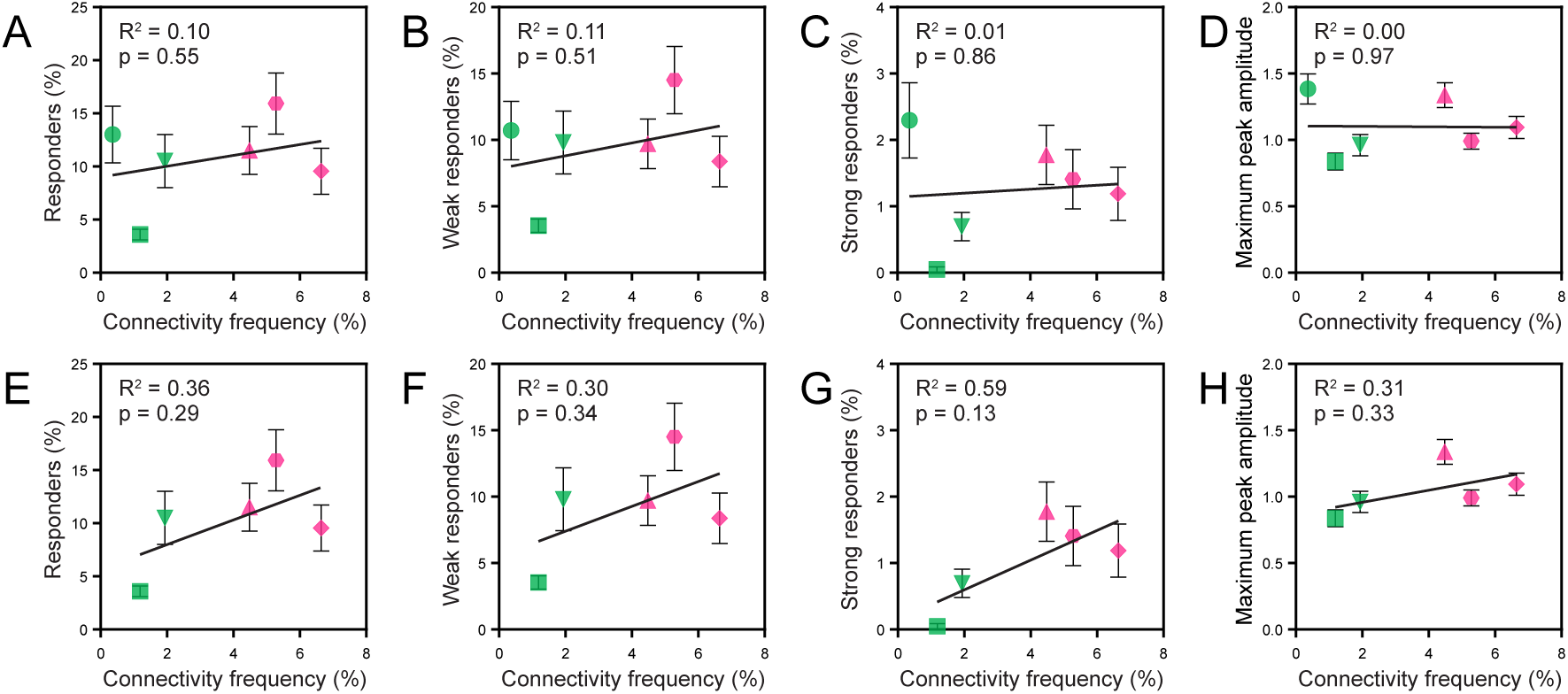
Optogenetic responses versus connectivity frequency from large-scale tracing datasets. Related to Figure 3 (A–H) Quantification of Kenyon cell responses plotted against connectivity frequency from large-scale tracing datasets. Panels A–D include all data; panels E–H show the same analyses with VL1 excluded. (A,E) Percentage of all responders. (B,F) Percentage of weak responders. (C,G) Percentage of strong responders. (D,H) Maximum peak ΔF/F amplitude. Each data point represents a *split-GAL4* line; error bars show standard error of the mean; linear regression lines are shown with R² and p values. Symbols indicate projection neuron identity: circle (VL1), square (DL3), downward triangle (VM4), upward triangle (DA1), hexagon (DA4l+DC3), diamond (VM5d+VM7d+VM7v). Colors indicate degree of representation: pink (overrepresented) and green (underrepresented).

**Figure S5.**
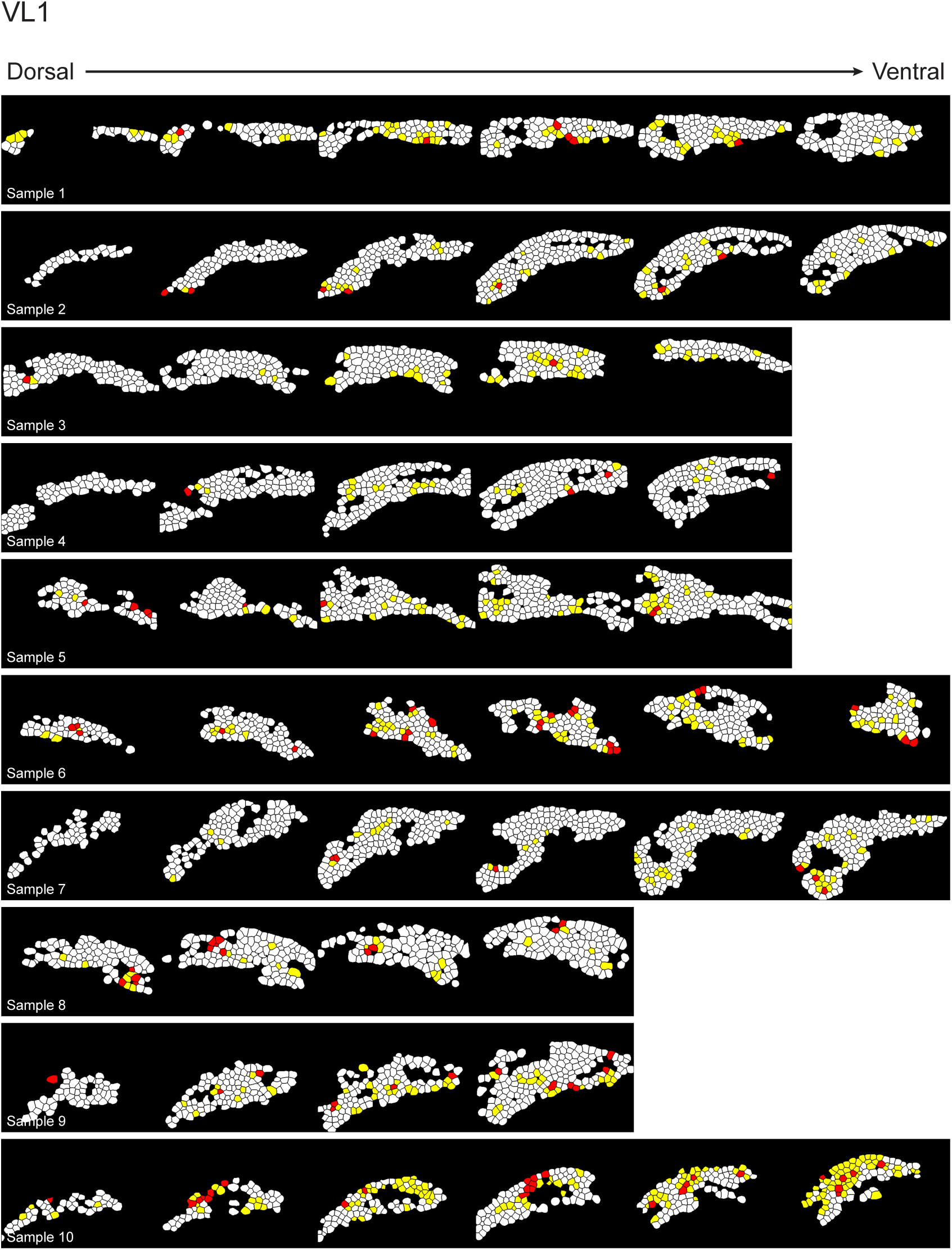
VL1 optogenetic responses across all samples. Related to Figure 3 Kenyon cell responses to VL1 optogenetic stimulation across ten samples. Each imaging plane are arranged from dorsal (left) to ventral (right), with a total of 5,015 cells recorded. Cells are classified as non-responders (white), weak responders (yellow), or strong responders (red) according to the criteria shown in Figure S2.

**Figure S6.**
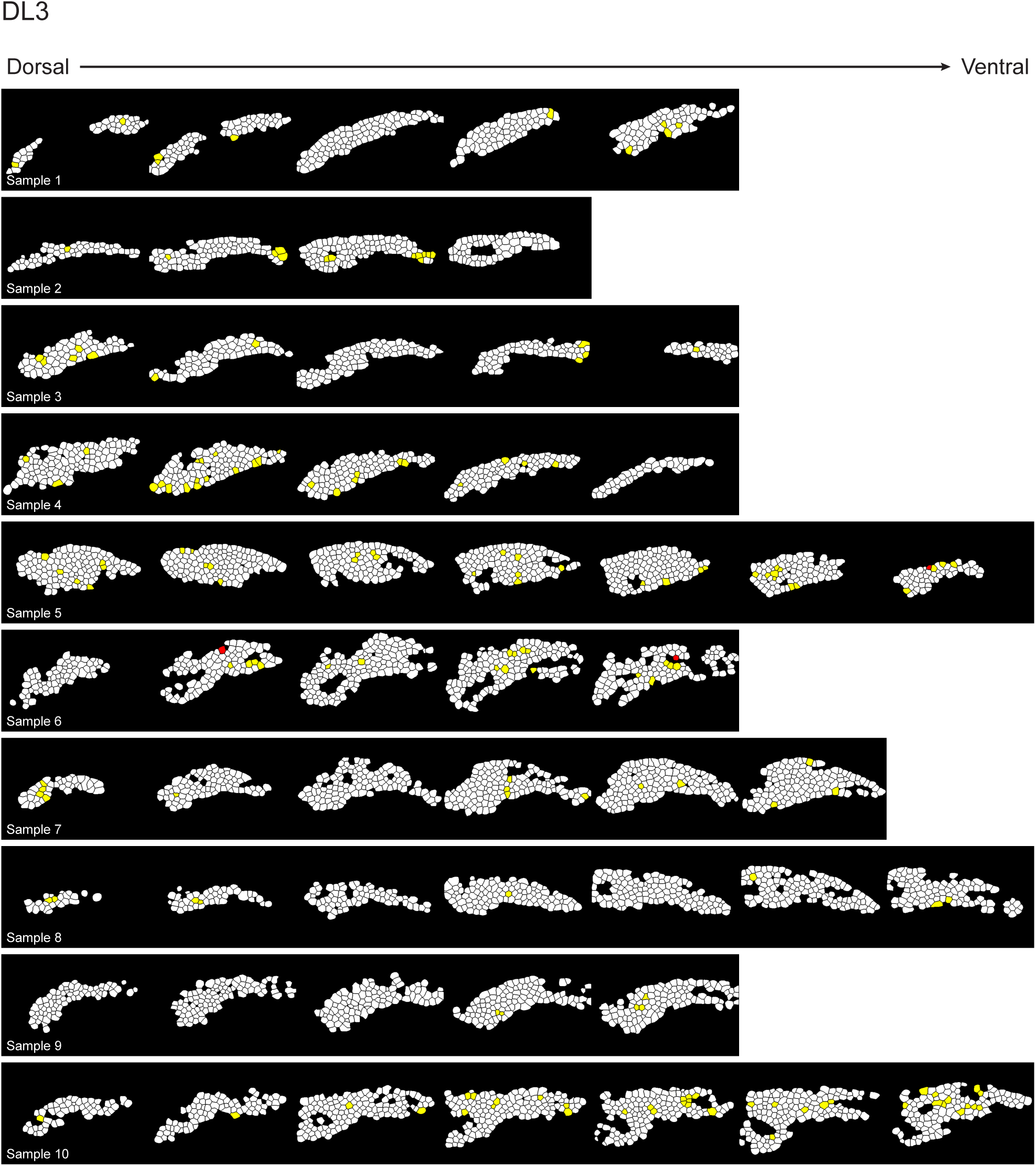
DL3 optogenetic responses across all samples. Related to Figure 3 Kenyon cell responses to DL3 optogenetic stimulation across ten samples. Each imaging plane are arranged from dorsal (left) to ventral (right), with a total of 5,059 cells recorded. Cells are classified as non-responders (white), weak responders (yellow), or strong responders (red) according to the criteria shown in Figure S2.

**Figure S7.**
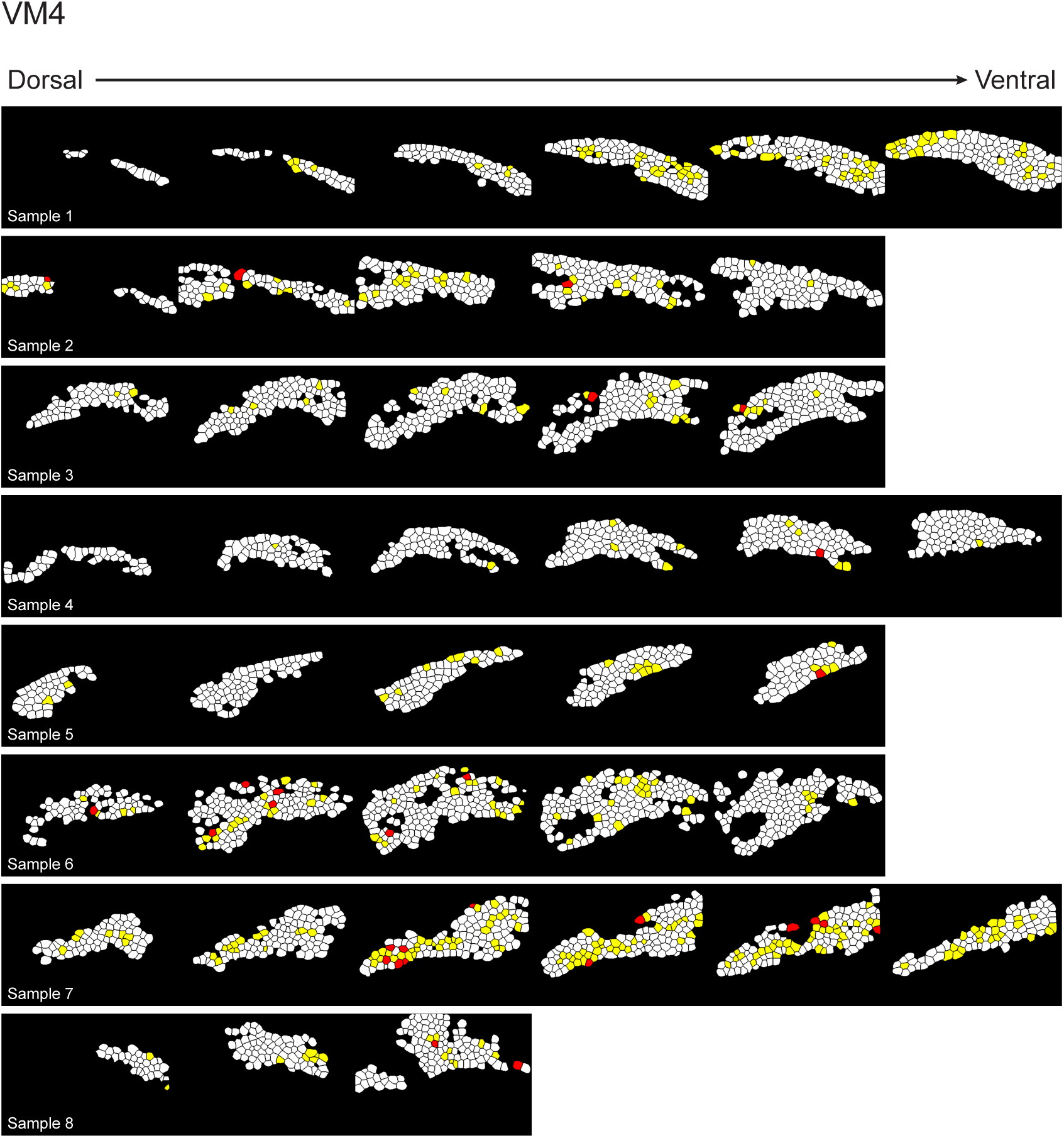
VM4 optogenetic responses across all samples. Related to Figure 3 Kenyon cell responses to VM4 optogenetic stimulation across eight samples. Each imaging plane are arranged from dorsal (left) to ventral (right), with a total of 3,590 cells recorded. Cells are classified as non-responders (white), weak responders (yellow), or strong responders (red) according to the criteria shown in Figure S2.

**Figure S8.**
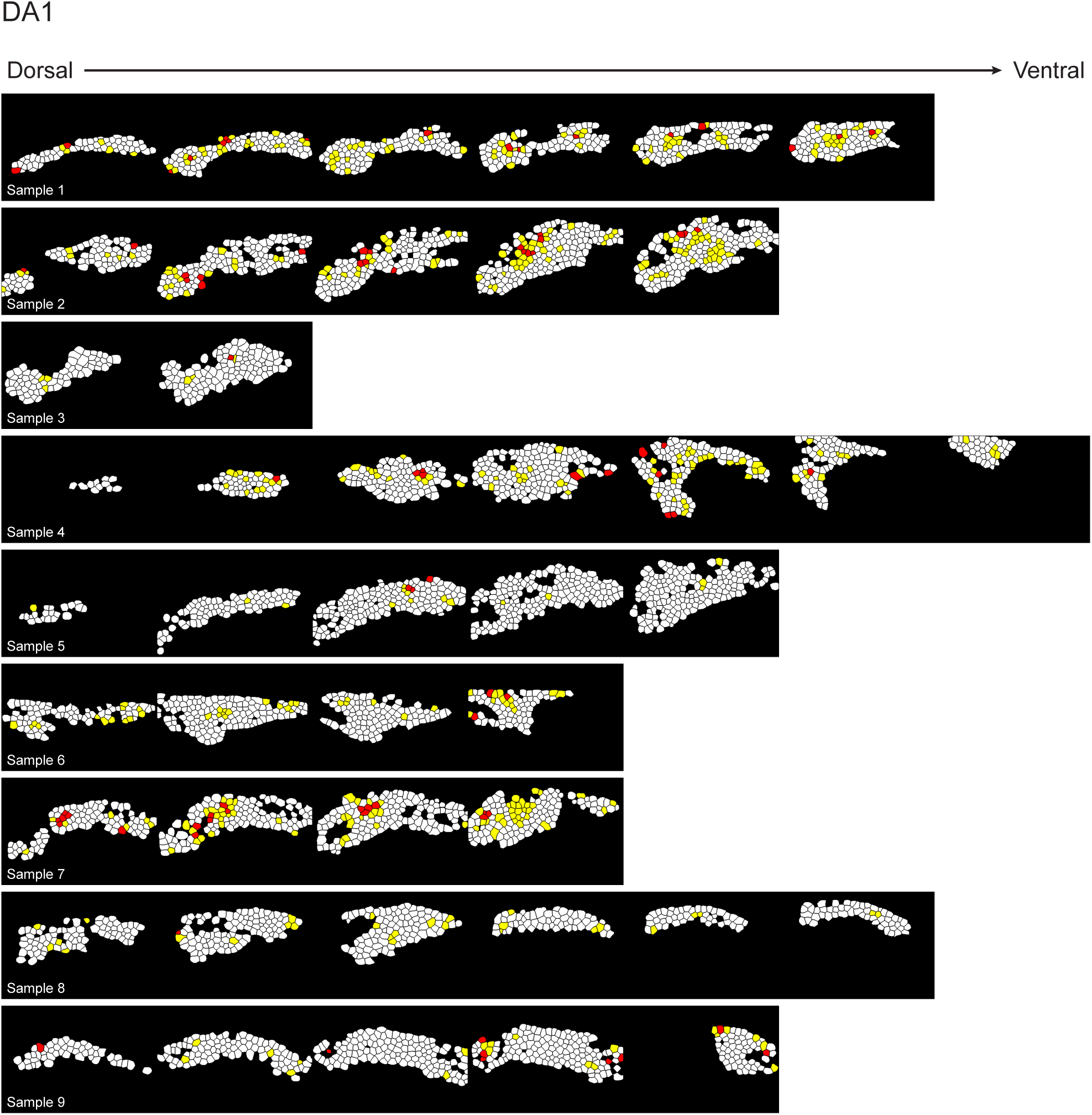
DA1 optogenetic responses across all samples. Related to Figure 3 Kenyon cell responses to DA1 optogenetic stimulation across nine samples. Each imaging plane are arranged from dorsal (left) to ventral (right), with a total of 4,432 cells recorded. Cells are classified as non-responders (white), weak responders (yellow), or strong responders (red) according to the criteria shown in Figure S2.

**Figure S9.**
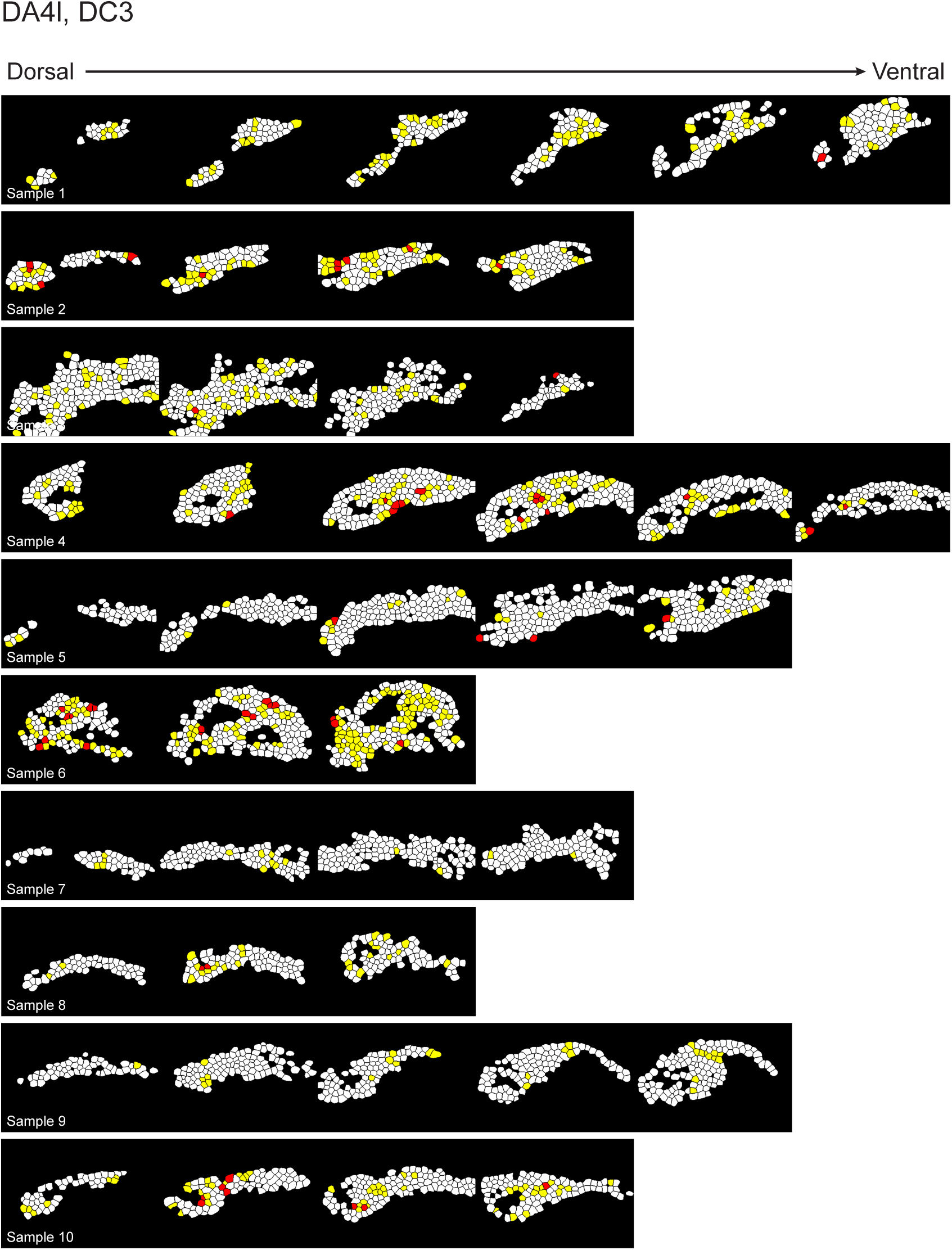
DA4l+DC3 optogenetic responses across all samples. Related to Figure 3 Kenyon cell responses to DA4l+DC3 optogenetic stimulation across ten samples. Each imaging plane are arranged from dorsal (left) to ventral (right), with a total of 4,199 cells recorded. Cells are classified as non-responders (white), weak responders (yellow), or strong responders (red) according to the criteria shown in Figure S2.

**Figure S10.**
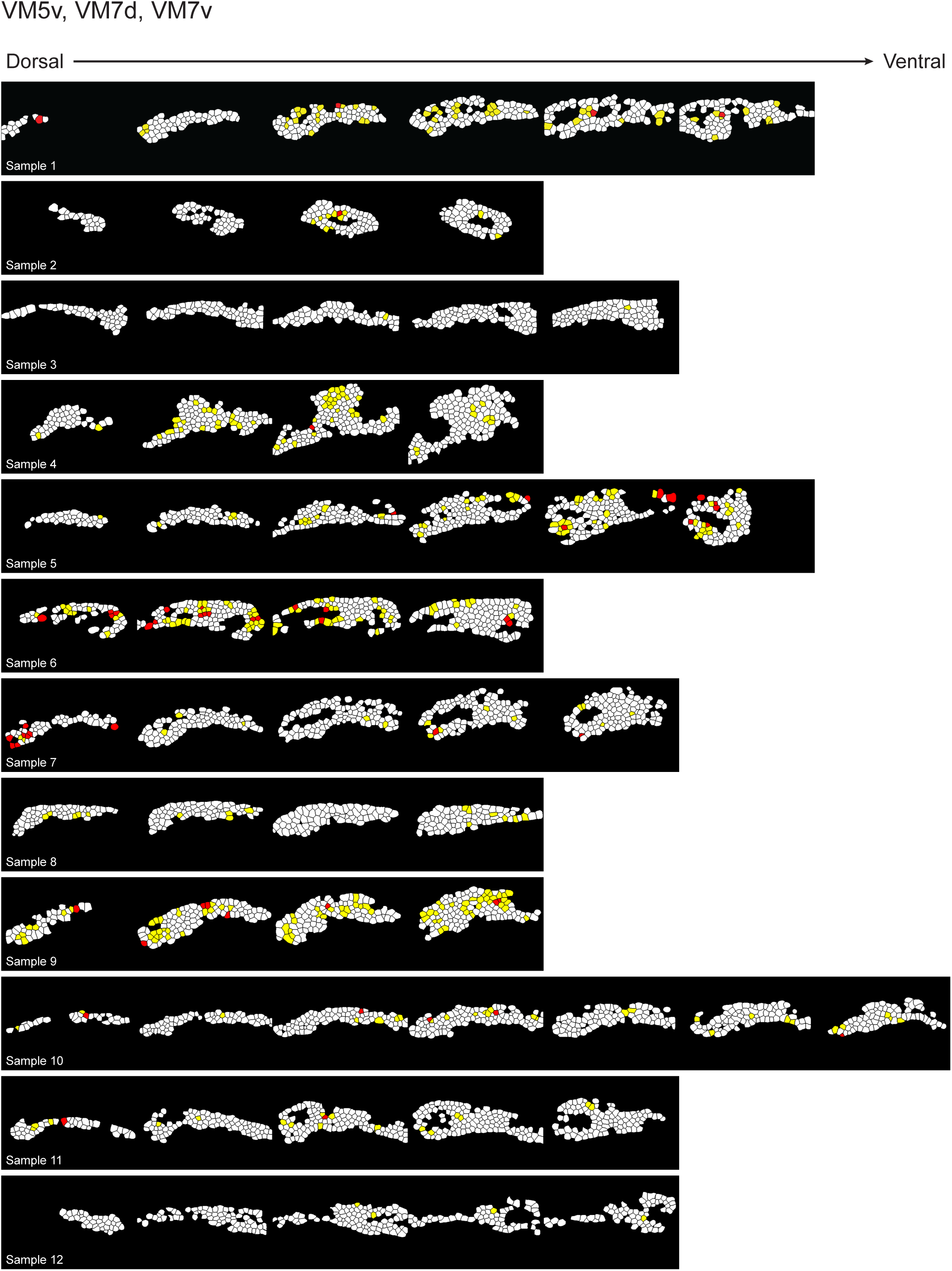
VM5d+VM7d+VM7v optogenetic responses across all samples. Related to Figure 3 Kenyon cell responses to VM5d+VM7d+VM7v optogenetic stimulation across twelve samples. Each imaging plane are arranged from dorsal (left) to ventral (right), with a total of 4,617 cells recorded. Cells are classified as non-responders (white), weak responders (yellow), or strong responders (red) according to the criteria shown in Figure S2.

**Figure S11.**
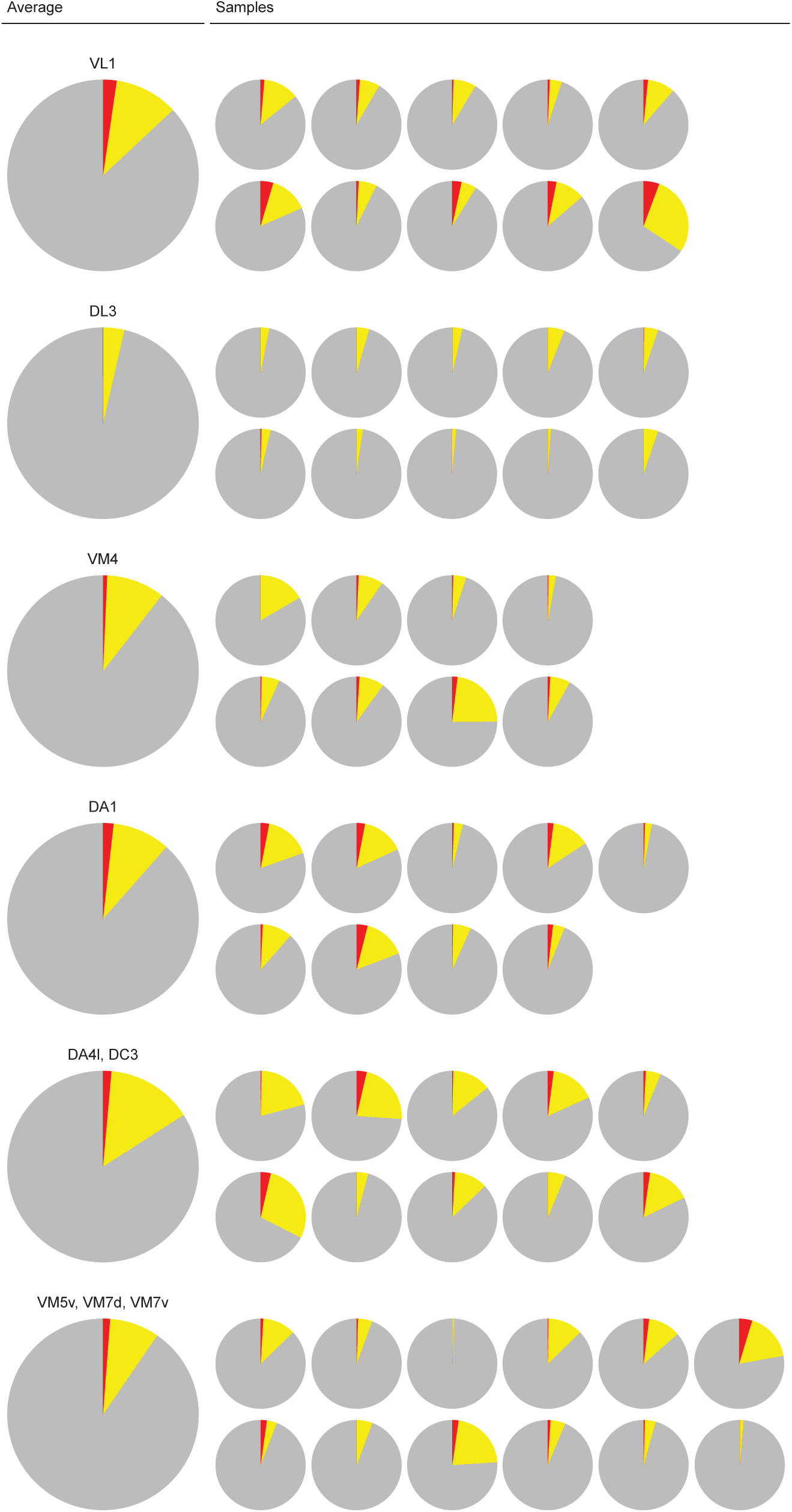
Response proportions for optogenetic experiments. Related to Figure 3 Pie charts showing proportions of non-responders (gray), weak responders (yellow), and strong responders (red) for each *split-GAL4* line across individual samples or average all sample in a data set.

**Figure S12.**
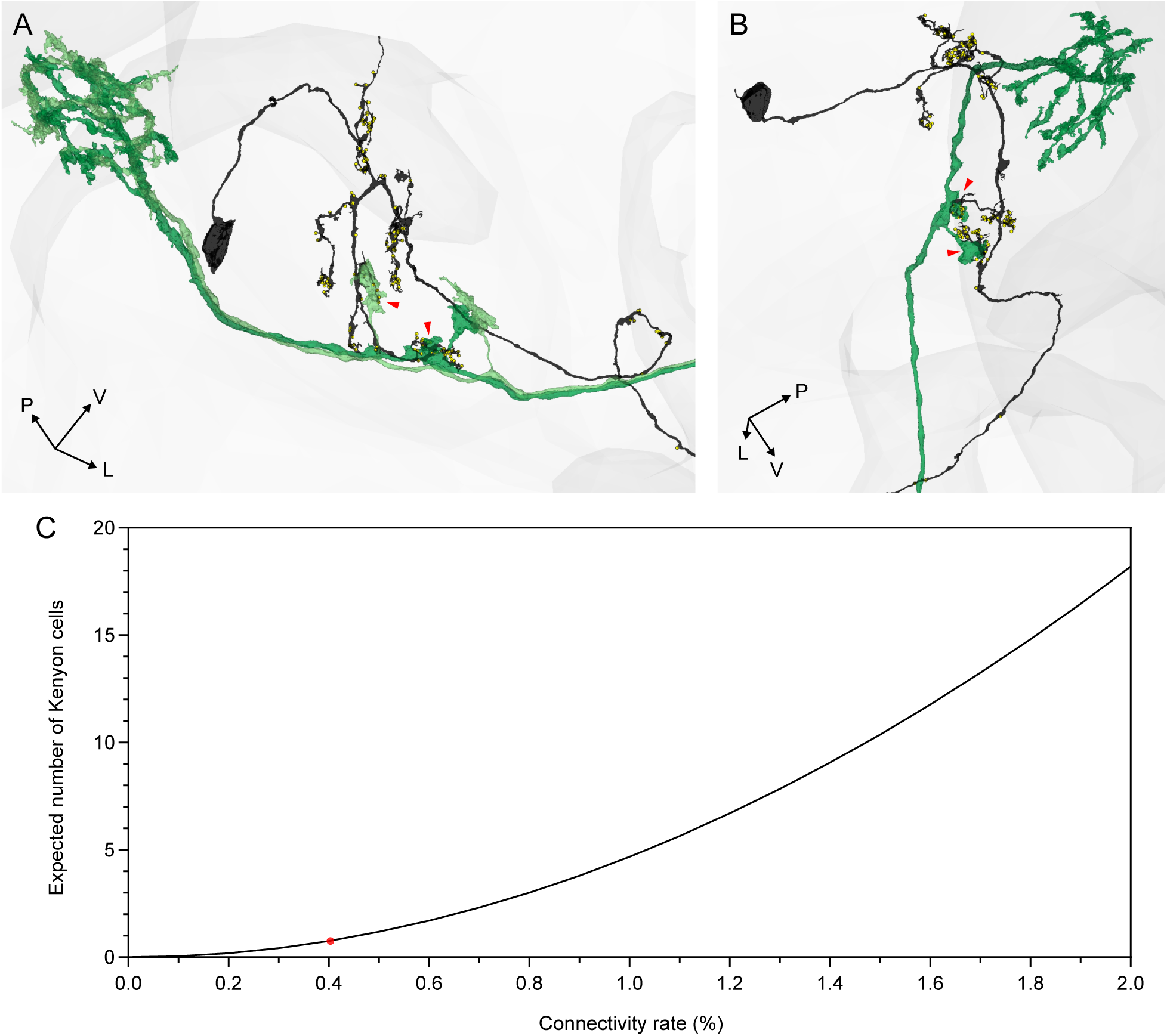
VL1 convergence analysis. Related to Figure 3 (A,B) Connectome reconstructions of the only two Kenyon cells receiving double VL1 inputs identified in the FlyWire connectome (Dorkenwald et al., 2024). (A) One Kenyon cell (black) receives inputs from two distinct VL1 projection neurons (light and dark green; red arrows). (B) Another Kenyon cell (black) receives two inputs from the same VL1 projection neuron (dark green). Orientation axes: P, posterior; V, ventral; L, lateral. (C) Probability of observing Kenyon cells with double inputs from the same glomerulus under a purely stochastic connectivity model, shown across different connectivity rates. The red marker indicates the connectivity frequency observed for VL1 in the connectome dataset (0.42%), which corresponds to one Kenyon cell with double VL1 inputs *per* mushroom body.

**Figure S13.**
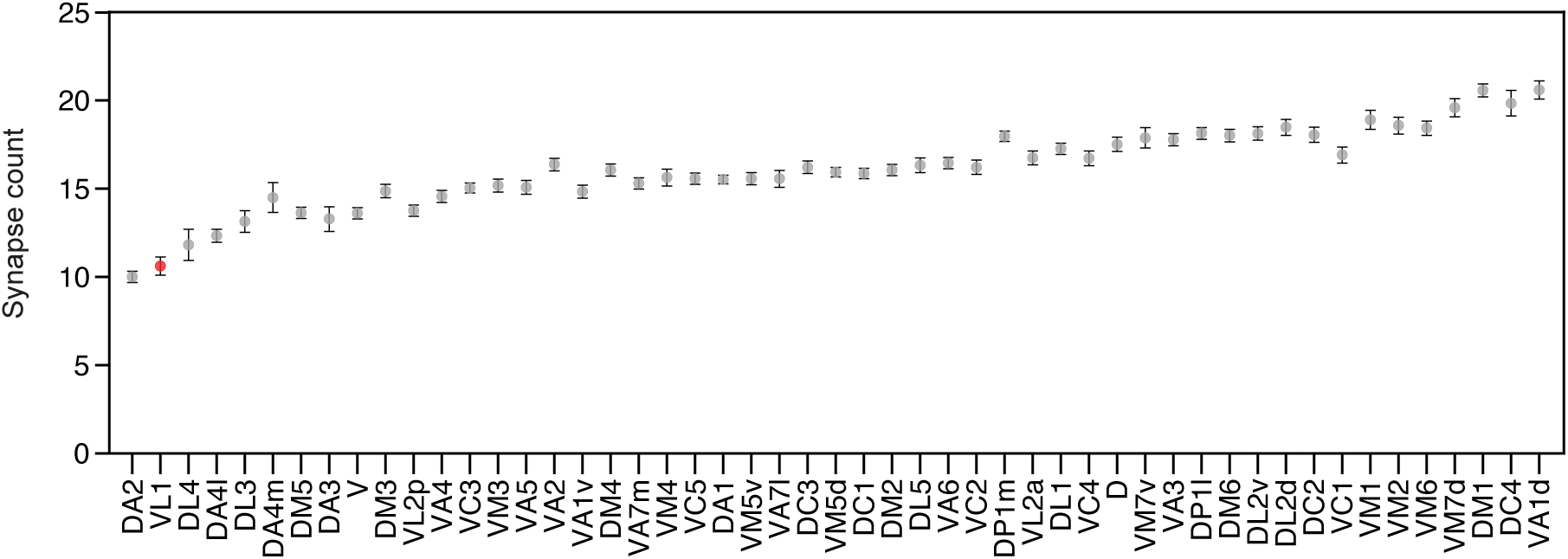
Synaptic strength across projection neuron types. Related to Figure 3 Number of synapses formed between Kenyon cells and projection neurons for each projection neuron type across all glomeruli identified in the FlyWire connectome (Dorkenwald et al., 2024). VL1 projection neurons are marked in red. Data points represent means; error bars show standard error of the mean.

**Figure S14.**
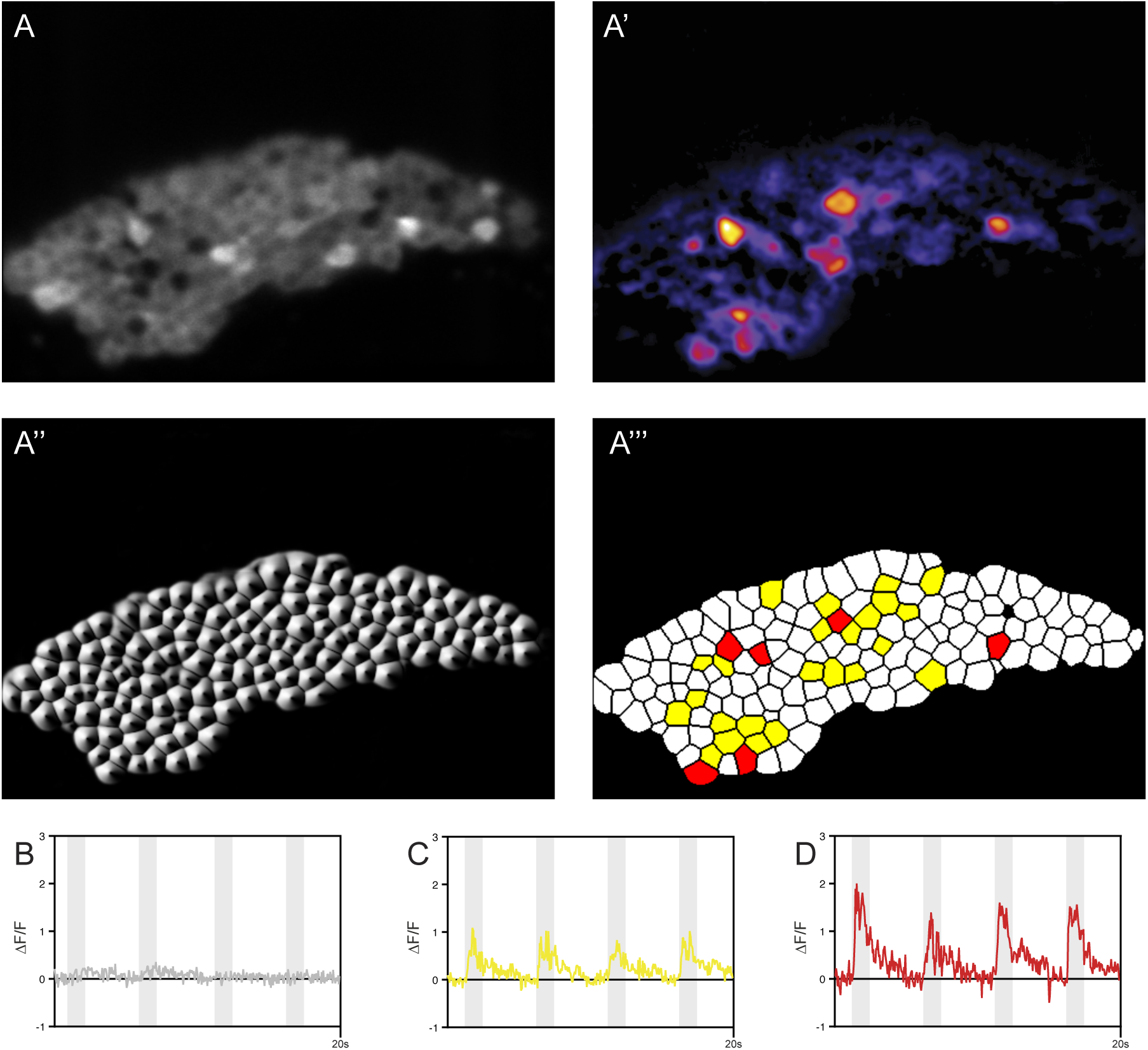
Classification of Kenyon cell responses to odor stimulation. Related to Figure 4 (A) Representative fluorescence image of Kenyon cells expressing GCaMP6f. (A’) Heat map of ΔF responses following odor presentation. (A’’) Segmented image showing individual Kenyon cell boundaries. (A’’’) Classification map showing non-responders (white), weak responders (yellow), and strong responders (red). (B–D) Representative calcium traces (ΔF/F) from non-responder (B), weak responder (C), and strong responder (D). Gray shaded boxes indicate three odor stimuli presentations per sample.

**Figure S15.**
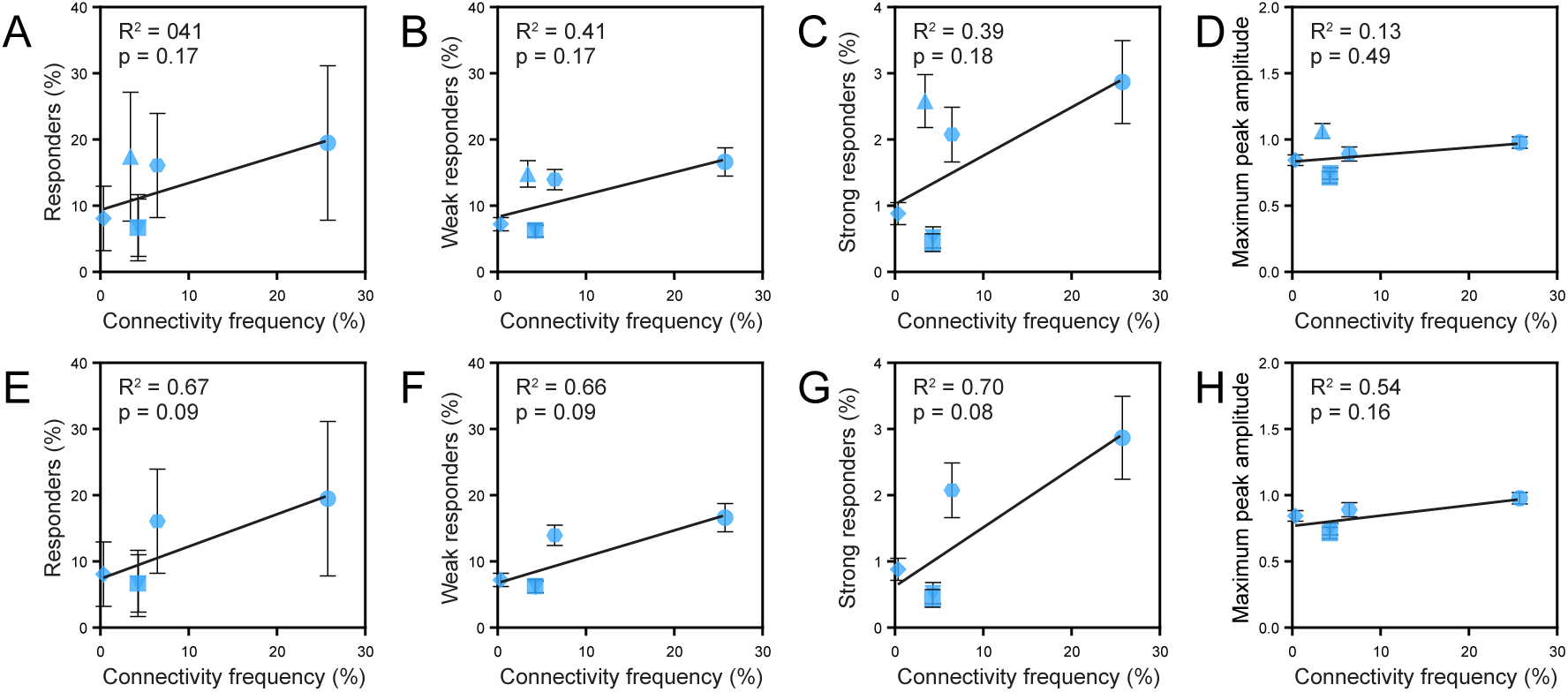
Odor responses versus connectivity frequency from large-scale tracing datasets. Related to Figure 4 (A–H) Quantification of Kenyon cell responses plotted against cumulative connectivity frequency from large-scale tracing datasets. Panels A–D include all odors; panels E–H show the same analyses with pyrrolidine excluded. (A,E) Percentage of all responders. (B,F) Percentage of weak responders. (C,G) Percentage of strong responders. (D,H) Mean maximum peak ΔF/F amplitude. Each data point represents an odor; error bars show standard error of the mean; linear regression lines are shown with R² and p values. Symbols indicate odor identity: square (farnesol), downward triangle (valencene), lozenge (geosmin), hexagon (4-methylcyclohexanol), upward triangle (pyrrolidine), circle (3-octanol).

**Figure S16.**
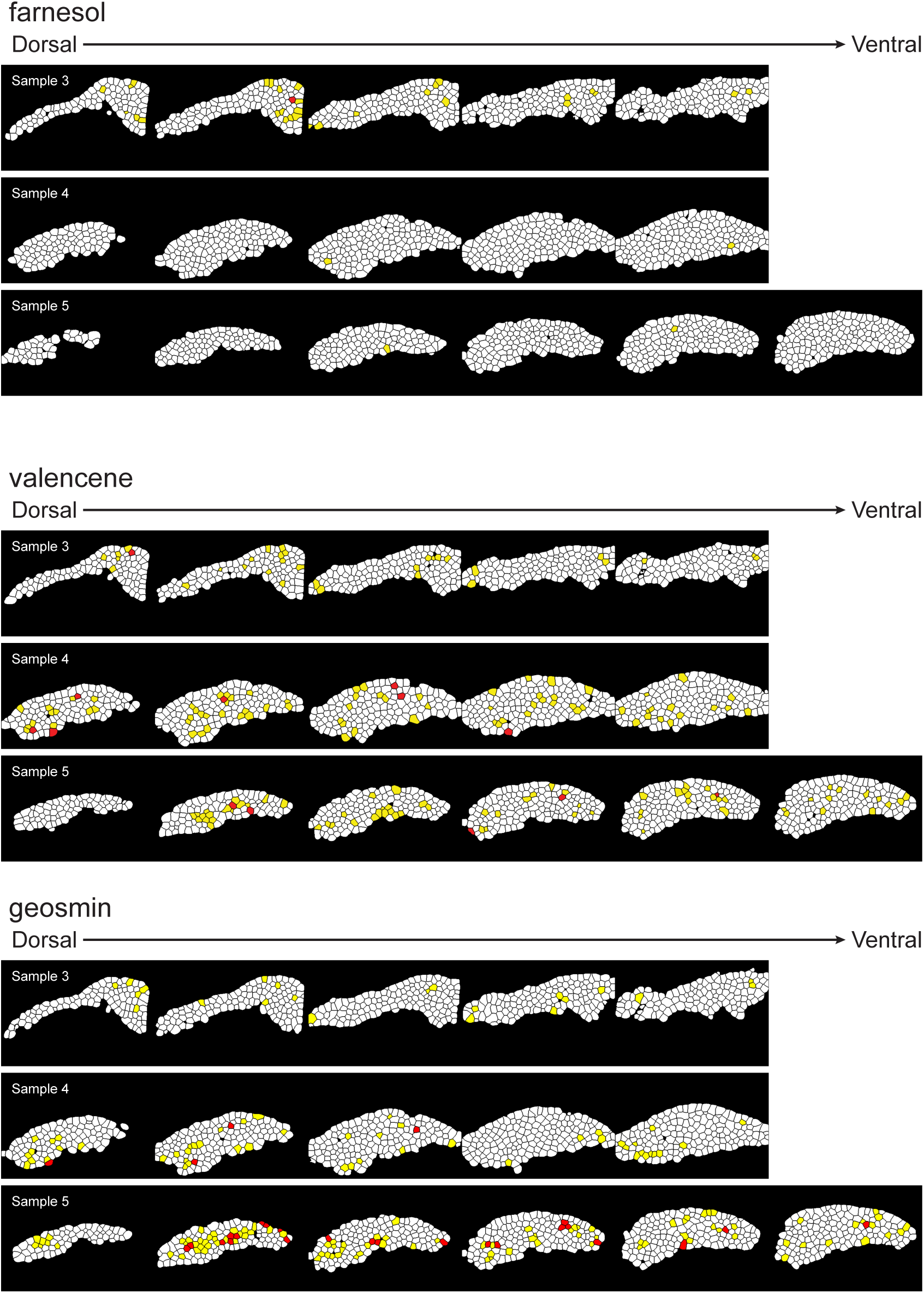
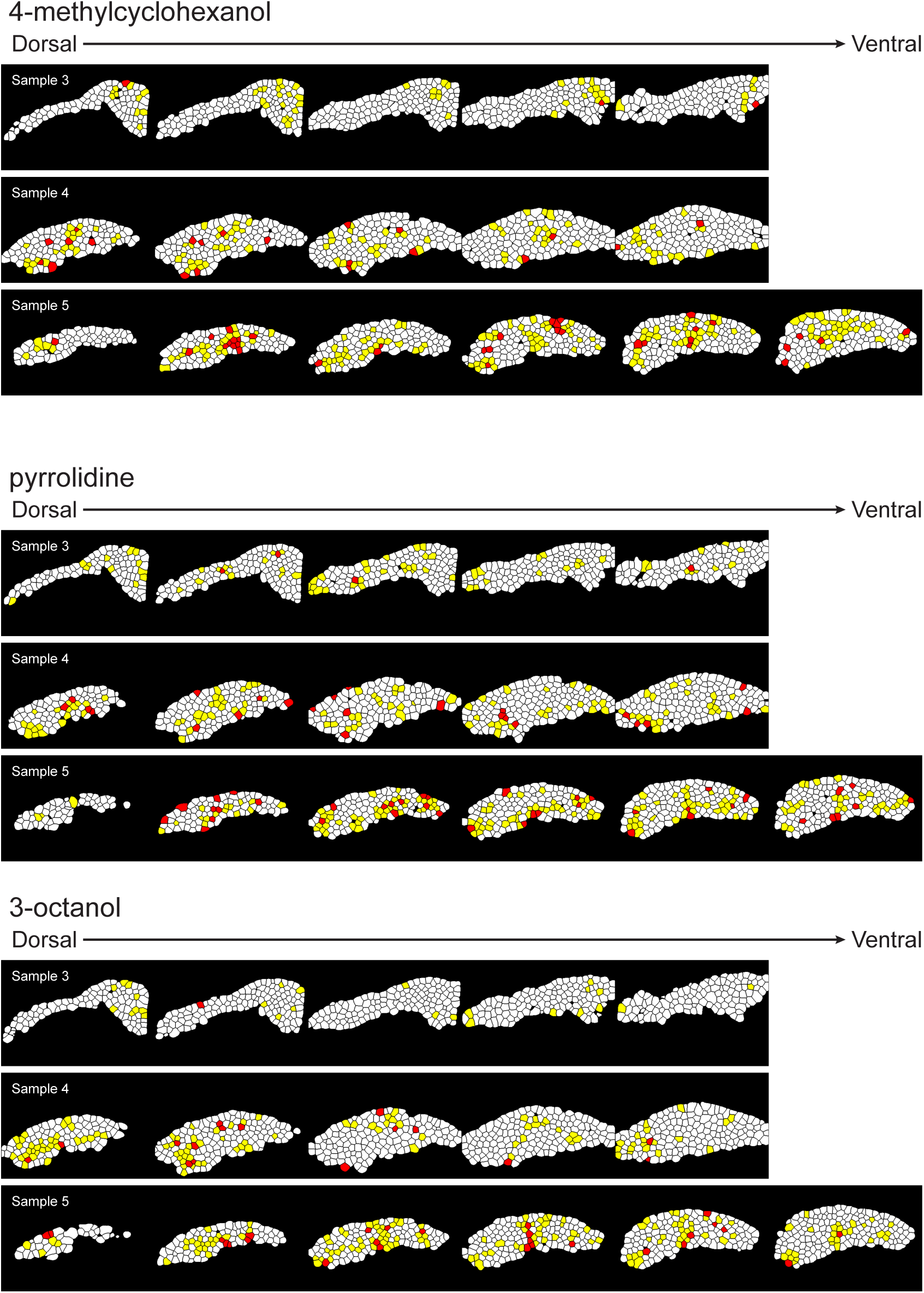
Odor-evoked responses across representative samples. Related to Figure 4 Kenyon cell responses to each odor across three representative samples. Each sample shows imaging planes from dorsal (left) to ventral (right). Cell counts: farnesol (617, 701 and 687 cells), valencene (602, 743 and 741 cells cells), geosmin (617, 727 and 777 cells), 4-methylcyclohexanol (617, 741 and 748 cells616), pyrrolidine (623, 685 and 723 cells), 3-octanol (597, 728 and 728 cells). Non-responders (white), weak responders (yellow), and strong responders (red) classified as in Figure S14.

**Figure S17.**
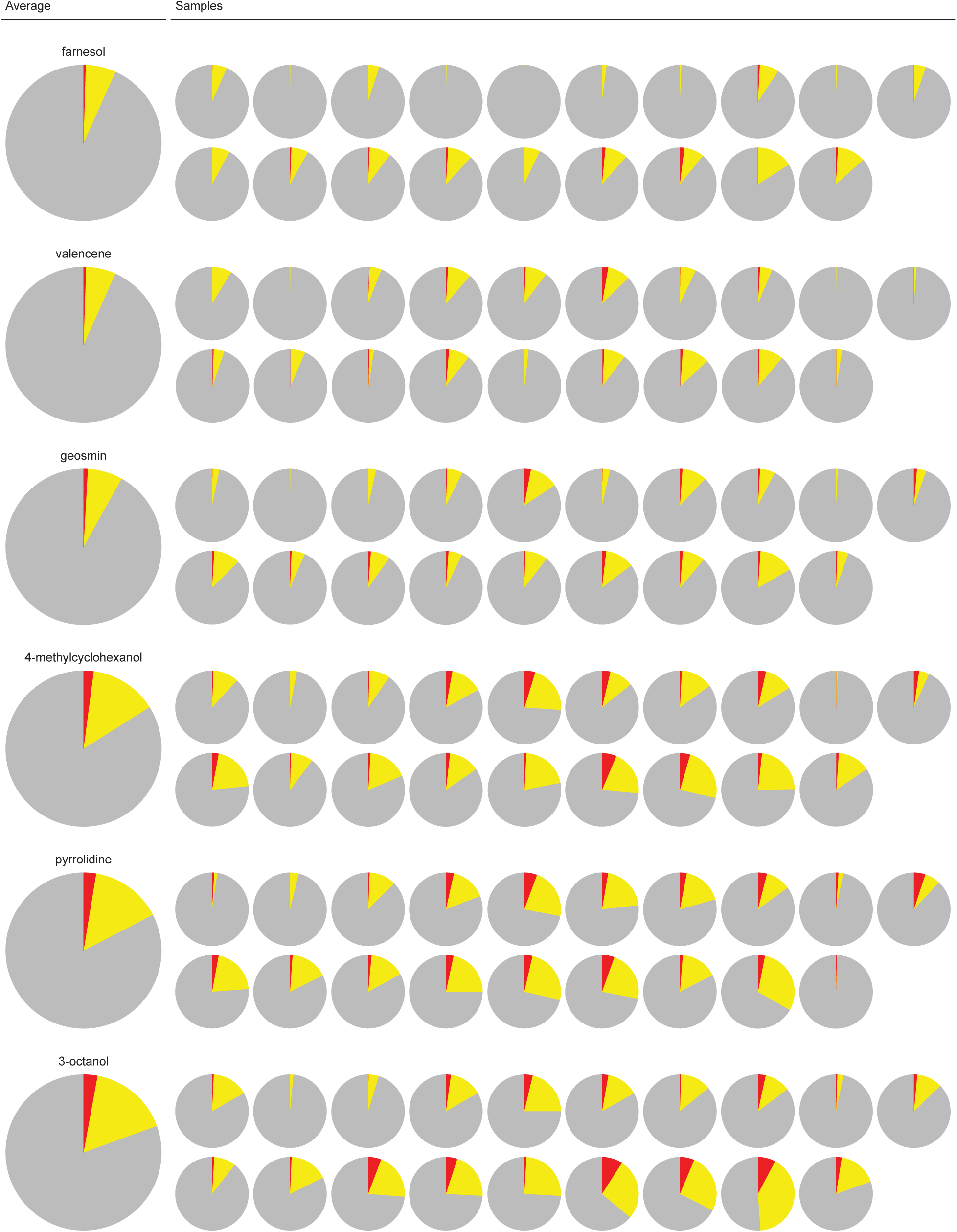
Response proportions for odor experiments. Related to Figure 4 Pie charts showing proportions of non-responders (gray), weak responders (yellow), and strong responders (red) for each odor across individual samples.

**Figure S18.**
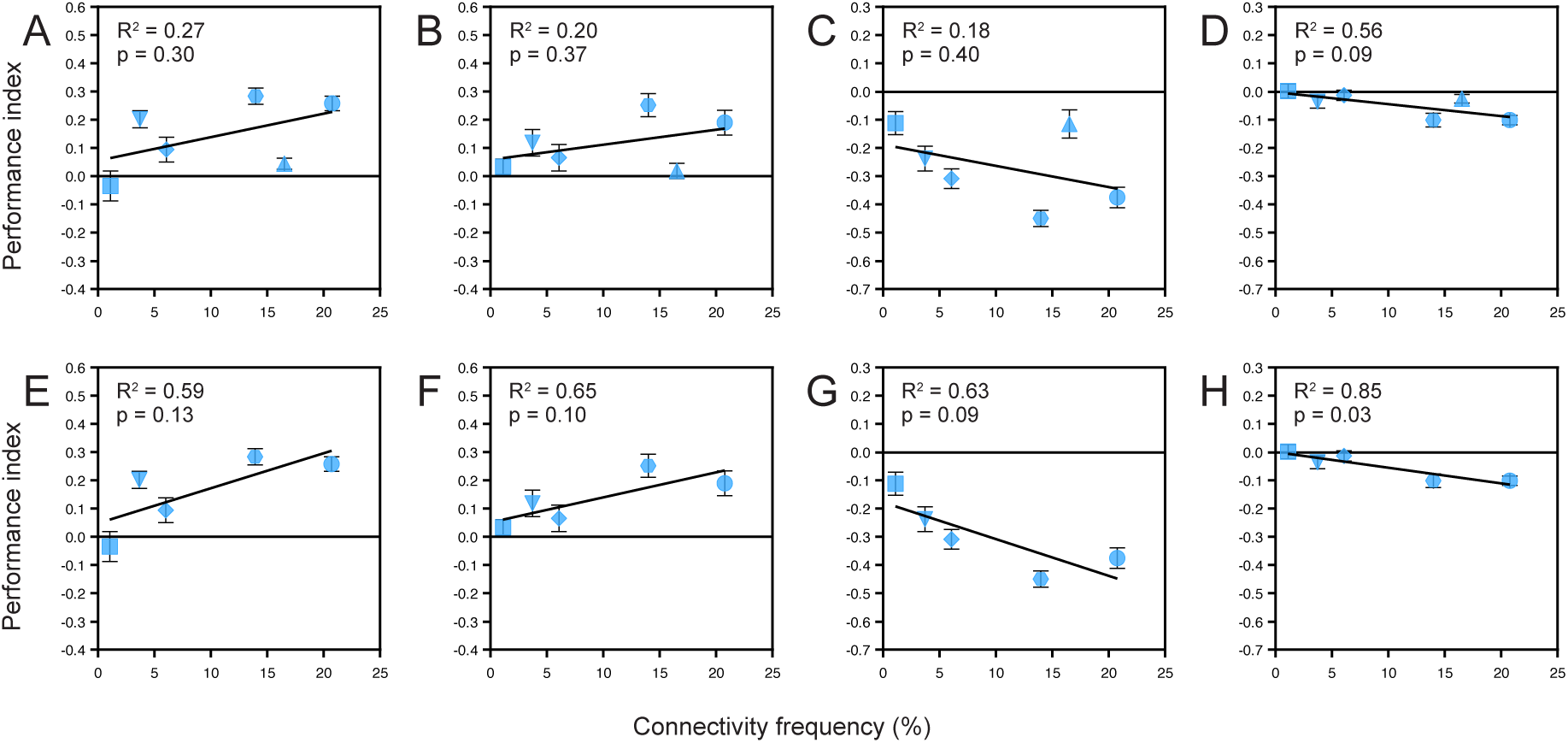
Learning performance versus connectivity frequency from large-scale tracing datasets. Related to Figure 5 (A–H) Correlation between learning performance and cumulative connectivity frequency from large-scale tracing datasets. Panels A–D include all odors; panels E–H show the same analyses with pyrrolidine excluded. Each data point represents an odor; error bars show standard error of the mean; linear regression lines are shown with R² and p values. Symbols indicate odor identity: square (farnesol), downward triangle (valencene), lozenge (geosmin), hexagon (4-methylcyclohexanol), upward triangle (pyrrolidine), circle (3-octanol).

## REFERENCES

Albus JS. 1971. A theory of cerebellar function. Mathematical Biosciencs, 10:25–61. DOI: 10.1016/0025-5564(71)90051-4

Amaral DG, Scharfman HE, Lavenex P. 2007. The dentate gyrus: Fundamental neuroanatomical organization (dentate gyrus for dummies) Progress in Brain Research 163:3–790 DOI: 10.1016/S0079-6123(07)63001-5, PMID: 17765709

Apps R, Hawkes R. 2009. Cerebellar cortical organization: A one-map hypothesis. Nature Reviews Neuroscience, 10:670–681. DOI: 10.1038/nrn2698, PMID: 19693030

Aso Y, Hattori D, Yu Y, Johnston RM, Iyer NA, Ngo TT, Dionne H, Abbott L, Axel R, Tanimoto H, Rubin GM. 2014. The neuronal architecture of the mushroom body provides a logic for associative learning. eLife, 3:e04577. DOI: 10.7554/eLife.04577, PMID: 25535793

Babadi B, Sompolinsky H. 2014. Sparseness and expansion in sensory representations. Neuron, 83:1213–1226. DOI: 10.1016/j.neuron.2014.07.035, PMID: 25155954

Bates AS, Schlegel P, Roberts RJV, Drummond N, Tamimi IFM, Turnbull R, Zhao X, Marin EC, Popovici PD, Dhawan S, Jamasb A, Javier A, Serratosa Capdevila L, Li F, Rubin GM, Waddell S, Bock DD, Costa M, Jefferis GSXE. 2020a. Complete connectomic reconstruction of olfactory projection neurons in the fly brain. Current Biology, 30:3183–3199.e6. DOI: 10.1016/j.cub.2020.06.042, PMID: 32619485

Bates AS, Manton JD, Jagannathan S, Costa M, Schlegel P, Rohlfing T, Jefferis GS. 2020b. The natverse, a versatile toolbox for combining and analysing neuroanatomical data. eLife, 9:e53350. DOI: 10.7554/eLife.53350, PMID: 32286229

Butcher NJ, Friedrich AB, Lu Z, Tanimoto H, Meinertzhagen IA. 2012. Different classes of input and output neurons reveal new features in microglomeruli of the adult *Drosophila* mushroom body calyx. Journal of Comparative Neurology, 520:2185–2201. DOI: 10.1002/cne.23037, PMID: 22237598

Caron SJC, Ruta V, Abbott LF, Axel R. 2013. Random convergence of olfactory inputs in the Drosophila mushroom body. Nature, 497:113–117. DOI: 10.1038/nature12063, PMID: 23615618

Cayco-Gajic NA, Silver RA. 2019. Re-evaluating circuit mechanisms underlying pattern separation. Neuron, 101:584–602. DOI: 10.1016/j.neuron.2019.01.044, PMID: 30790539

Cognigni P, Felsenberg J, Waddell S. 2018. Do the right thing: Neural network mechanisms of memory formation, expression and update in Drosophila. Current Opinion in Neurobiology, 49:51–58. DOI: 10.1016/j.conb.2017.12.002, PMID: 29258011

Dorkenwald S, Matsliah A, Sterling AR, Schlegel P, Yu S, McKellar CE, Lin A, Costa, M, Eichle K., Yin Y, Silversmith W, Schneider-Mizell C, Jordan CS, Brittain D, Halageri A, Kuehner K, Ogedengbe O, Morey R, Gager, J, Kruk K, Perlman E, Yang R, Deutsch D, Bland D, Sorek M, Lu R, Macrina T, Lee K, Bae, JA, Mu S, Nehoran B, Mitchell E, Popovych S, Wu J, Jia Z, Castro MA, Kemnitz N, Ih D, Bates AS, Eckstein N, Funke J, Collman F, Bock DD, Jefferis GSXE, Seung HS, Murthy M, FlyWire Consortium. 2024. Neuronal wiring diagram of an adult brain. Nature, 634:124–138. DOI: 10.1038/s41586-024-07558-y, PMID: 39358518

Ellis KE, Bervoet S., Smihula H, Ganguly I, Vigato E, Auer TO, Benton R, Litwin-Kumar A, Caron SJC. 2024. Evolution of connectivity architecture in the Drosophila mushroom body. Nature Communications, 15:4872. DOI: 10.1038/s41467-024-48839-4, PMID: 38849331

Ganguly I, Heckman EL, Litwin-Kumar A, Clowney EJ, Behnia R. 2024. Diversity of visual inputs to Kenyon cells of the Drosophila mushroom body. Nature Communications, 15:5698. DOI: 10.1038/s41467-024-49616-z, PMID: 37873086

Giovannucci A, Friedrich J, Gunn P, Kalfon J, Brown BL, Koay SA, Taxidis J, Najafi F, Gauthier JL, Zhou P, Khakh BS, Tank D W, Chklovskii DB, Pnevmatikakis EA. 2019. CaImAn an open source tool for scalable calcium imaging data analysis. eLife, 8:e38173. DOI: 10.7554/eLife.38173, PMID: 30652683

Grabe V, Strutz A, Baschwitz A, Hansson B, Sachse S. 2015. Digital *in vivo* 3D atlas of the antennal lobe of *Drosophila melanogaster*. Journal of Comparative Neurology, 523:530–544. 10.1002/cne.23697, PMID: 25327641

Hallem EA, Carlson JR. 2006. Coding of odors by a receptor repertoire. Cell, 125:143–160. DOI: 10.1016/j.cell.2006.01.050, PMID: 16615896

Hargreaves EL, Rao G, Lee I, Knierim JJ. 2005. Major dissociation between medial and lateral entorhinal input to dorsal hippocampus. Science, 308:1792–1794. DOI: 10.1126/science.1110449, PMID: 15961670

Hartmann T, Ober D. 2008. Defense by pyrrolizidine alkaloids: Developed by plants and recruited by insects. Induced Plant Resistance to Herbivory. DOI: 10.1007/978-1-4020-8182-8_10, ISBN: 978-1-4020-8181-1

Hayashi TT, MacKenzie AJ, Ganguly I, Ellis KE, Smihula HM, Jacob MS, Litwin-Kumar A, Caron SJC. 2022. Mushroom body input connections form independently of sensory activity in Drosophila melanogaster. Current Biology, 32:4000–4012.e5. 10.1016/j.cub.2022.07.055, PMID: 35977547

Hunter JD. 2007. Matplotlib: A 2D graphics environment. Computing in Science & Engineering, 9:90–95. DOI: 10.1109/MCSE.2007.55, ISSN: 1521-9615

Knierim JJ, Neunuebel JP, Deshmukh SS. 2014. Functional correlates of the lateral and medial entorhinal cortex: Objects, path integration and local–global reference frames. Philosophical Transactions of the Royal Society B: Biological Sciences, 369:20130369. DOI: 10.1098/rstb.2013.0369, PMID: 24366146

Krashes M, Waddell S. 2011a. Drosophila appetitive olfactory conditioning. Cold Spring Harbor Protocols, 5. DOI: 10.1101/pdb.prot5609, PMID: 21536767

Krashes M, Waddell S. 2011b. Drosophila aversive olfactory conditioning. Cold Spring Harbor Protocols, 5. DOI: 10.1101/pdb.prot5608, PMID: 21536768

Li F, Lindsey J., Marin EC, Otto N, Dreher M, Dempsey G, Stark I, Bates AS, Pleijzier, MW, Schlegel P, Nern A, Takemura S, Eckstein N, Yang T, Francis A, Braun A, Parekh, R, Costa M, Scheffer LK, Aso Y, Jefferis GSXE, Abbot LF, Litwin-Kumar A, Waddell S, Rubin GM. 2020. The connectome of the adult Drosophila mushroom body provides insights into function. eLife, 9:e62576. DOI: 10.7554/eLife.62576, PMID: 33315010

Litwin-Kumar A, Harris KD, Axel R, Sompolinsky H, Abbott LF. 2017. Optimal degrees of synaptic connectivity. Neuron, 93:1153–1164.e7. DOI: 10.1016/j.neuron.2017.01.030, PMID: 28215558

Marr, D. 1969. A theory of cerebellar cortex. The Journal of Physiology, 202:437–470. DOI: 10.1113/jphysiol.1969.sp008820, PMID: 5784296

Münch D, Galizia CG. 2016. DoOR 2.0—Comprehensive mapping of Drosophila melanogaster odorant responses. Scientific Reports, 6: 21841. DOI: 10.1038/srep21841, PMID: 26912260

Nguyen TM, Thomas LA, Rhoades JL, Ricchi I, Yuan XC, Sheridan A, Hildebrand DGC, Funke J, Regehr WG, Lee WCA. 2023. Structured cerebellar connectivity supports resilient pattern separation. Nature, 613:543–549. DOI: 10.1038/s41586-022-05471-w, PMID: 36418404

Olsen SR, Wilson RI. 2008. Lateral presynaptic inhibition mediates gain control in an olfactory circuit. Nature, 452:956–960. DOI: 10.1038/nature06864, PMID: 18344978

Pedregosa F, Varoquaux, G, Org N, Gramfort, A, Michel V, Fr L, Thirion B, Grisel O, Blondel M, Prettenhofer P, Weiss R, Dubourg V, Vanderplas J, Passos A, Cournapeau, D. 2011. Scikit-learn: Machine learning in python. Journal of Machine Learning Research, 12:2825–2830

Prieto-Godino LL, Rytz R, Cruchet S, Bargeton B, Abuin L, Silbering AF, Ruta V, Dal Peraro M, Benton R. 2017. Evolution of acid-sensing olfactory circuits in Drosophilids. Neuron, 93:661–676.e6. DOI: 10.1016/j.neuron.2016.12.024, PMID: 28111079

Scheffer LK, Xu CS, Januszewski M, Lu Z, Takemura S, Hayworth KJ, Huang GB, Shinomiya K, Maitlin-Shepard J, Berg S, Clements J, Hubbard PM, Katz WT, Umayam L, Zhao T, Ackerman D, Blakely T, Bogovic J, Dolafi T, … Plaza SM. 2020. A connectome and analysis of the adult Drosophila central brain. eLife, 9:e57443. DOI: 10.7554/eLife.57443, PMID: 32880371

Schindelin J, Arganda-Carreras I, Frise E, Kaynig V, Longair M, Pietzsch T, Preibisch S, Rueden C, Saalfeld S, Schmid B, Tinevez JY, White DJ, Hartenstein V, Eliceiri K, Tomancak P, Cardona A. 2012. Fiji: An open-source platform for biological-image analysis. Nature Methods, 9:676–682. DOI: 10.1038/nmeth.2019, PMID: 22743772

Shuai Y, Sammons M, Sterne GR, Hibbard KL, Yang H, Yang CP, Managan C, Siwanowicz I, Lee T, Rubin GM, Turner GC, Aso Y. 2025. Driver lines for studying associative learning in Drosophila. eLife, 13:RP94168. DOI: 10.7554/eLife.94168, PMID: 39879130

Silbering AF, Rytz R, Grosjean Y, Abuin L, Ramdya P, Jefferis GSXE, Benton R. 2011. Complementary function and integrated wiring of the evolutionarily distinct *Drosophila* olfactory subsystems. The Journal of Neuroscience, 31:13357–13375. DOI: 10.1523/JNEUROSCI.2360-11.2011, PMID: 21940430

Stensmyr MC, Dweck HKM, Farhan A, Ibba I, Strutz A, Mukunda L, Linz J, Grabe V, Steck K, Lavista-Llanos S, Wicher D, Sachse S, Knaden M, Becher PG, Seki Y, Hansson BS. 2012. A conserved dedicated olfactory circuit for detecting harmful microbes in Drosophila. Cell, 151:1345–1357. DOI: 10.1016/j.cell.2012.09.046, PMID: 23217715

Stringer C, Wang T, Michaelos M, Pachitariu M. 2021. Cellpose: A generalist algorithm for cellular segmentation. Nature Methods, 18:100–106. DOI: 10.1038/s41592-020-01018-x, PMID: 33318659

Wan Y, Otsuna H, Holman HA, Bagley B, Ito M, Lewis AK, Colasanto M, Kardon G, Ito K, Hansen C. 2017. FluoRender: Joint freehand segmentation and visualization for many-channel fluorescence data analysis. BMC Bioinformatics, 18:280. DOI: 10.1186/s12859-017-1694-9, PMID: 28549411

Wilson RI. 2013. Early olfactory processing in *Drosophila*: Mechanisms and principles. Annual Review of Neuroscience, 36:217–241. DOI: 10.1146/annurev-neuro-062111-150533, PMID: 23841839

Xia R, Chen X, Engel TA, Moore T. 2024. Common and distinct neural mechanisms of attention. Trends in Cognitive Sciences, 28:554–567. DOI: 10.1016/j.tics.2024.01.005, PMID: 38388258

Zavitz D, Amematsro EA, Borisyuk A, Caron, SJC. 2021. Connectivity patterns that shape olfactory representation in a mushroom body network model. BioRxiv, DOI: 10.1101/2021.02.10.430647

Zheng Z, Li F, Fisher C, Ali IJ, Sharifi N, Calle-Schuler S, Hsu J, Masoodpanah N, Kmecova L, Kazimiers T, Perlman E, Nichols M, Li PH, Jain V, Bock DD. 2022. Structured sampling of olfactory input by the fly mushroom body. Current Biology, 32:3334–3349.e6. DOI: 10.1016/j.cub.2022.06.031, PMID: 35797998

